# Whole-brain mapping of TMS-induced discomfort with large-scale concurrent TMS-fMRI

**DOI:** 10.64898/2026.02.27.708526

**Authors:** Zhuoran Li, Yong Jiao, Yu Zhang, Nandi Zhang, Amit Etkin, Aaron D. Boes, Desmond J. Oathes, Jing Jiang

## Abstract

Transcranial magnetic stimulation (TMS) is a cornerstone tool for causal inference in human brain function and an increasingly used neuromodulation therapy, yet it induces well-recognized discomfort that may systematically bias measured outcomes. Despite its ubiquity, a critical gap remains in understanding how TMS-induced discomfort is represented across the brain and to what extent it contributes to TMS-evoked neural responses. Using concurrent TMS-fMRI across 11 cortical targets, we collected an unprecedented dataset (165 participants; 1,535 runs) spanning healthy participants and those with elevated affective symptoms. Cross-validated multivariate modeling revealed that TMS-induced discomfort engages distributed cortical and subcortical regions across sensorimotor, attentional, default mode, and limbic networks, with both shared and group-specific patterns. Discomfort-related activity accounted for approximately 12% and 25% of TMS-evoked responses in healthy and elevated-symptom groups, respectively, and varied systematically across stimulating sites. These findings identify TMS-induced discomfort as a substantial and previously under-characterized component of TMS-evoked neural responses, underscoring the need to explicitly measure and model it. By providing a whole-brain map of regional contributions associated with TMS-induced discomfort and an analytic framework to dissociate direct neuromodulatory effects from discomfort-related responses, this work improves the interpretability of TMS-evoked signals and supports more rigorous causal inference and therapeutic applications.

## Introduction

Since its development in 1985 (1), transcranial magnetic stimulation (TMS) has become a versatile brain stimulation tool for causal inference of brain functions and advancing noninvasive neuromodulation-based clinical therapies (2, 3). TMS delivers brief, high-intensity magnetic pulses that penetrate the scalp and induce activity changes in targeted cortical tissue beneath the coil. These pulses also stimulate scalp nociceptive fibers, muscles, and surrounding tissues, producing discomfort or pain that varies across scalp locations (4) and individuals (5). Although typically transient, such discomfort can increase psychological burden and contribute to dropout in both experimental research and clinical treatment (6, 7). Moreover, commonly used stimulation targets, particularly frontal regions, often induce stronger discomfort that has been shown to influence task performance (4, 8, 9), suggesting engagement of additional neurocognitive processes beyond basic sensory perception. This introduces a potential confound that can bias measured outcomes, complicate interpretations, and undermine causal inferences from TMS studies (10). Despite its widespread recognition and potential impact, no study has systematically mapped the neural correlates of TMS-induced discomfort, a critical step toward disentangling direct neuromodulatory effects from discomfort-driven influences and for optimizing TMS use in research and clinical practice.

Addressing this gap requires whole-brain characterization of neural responses to TMS-induced discomfort, as this experience likely engages a constellation of sensory, nociceptive, attentional, and affective processes distributed across the brain. TMS-induced discomfort or pain arises most likely from peripheral sensations of stimulation of trigeminal nerves, muscles, and/or nociceptors in the scalp beneath the coil or surrounding area. Therefore, it is expected to drive tactile sensory responses in primary and secondary somatosensory cortices (11) and recruit neural circuits involved in nociceptive processing (12) and pain (13). Moreover, the abrupt, high-salience nature of stimulation pulses may engage insula and anterior cingulate regions implicated in arousal, stimulus evaluation, and threat appraisal (14). Behavioral evidence further indicates that stronger TMS-induced discomfort is associated with slower responses and poorer task performance (4, 8, 9), suggesting shifts in attention and potential engagement of attentional networks (14, 15) and other higher-order systems (16, 17) involved in anticipatory, emotion regulation, and cognitive control processes. Given the distributed nature of these putative neural processes, TMS concurrent with functional magnetic resonance imaging (concurrent TMS-fMRI) (18, 19) presents a powerful approach to capture whole-brain responses associated with TMS-induced discomfort.

Several other gaps in the existing literature also limit our understanding of TMS-induced discomfort. First, most prior studies have examined only one or two stimulation sites. Because TMS-induced discomfort varies substantially across scalp locations (4), limited site sampling constrains the range of discomfort variability needed to map its neural correlates and disentangle discomfort-related neural effects from site-specific TMS neuromodulatory effects. Incorporating a broader set of commonly targeted regions is therefore crucial for determining core brain regions consistently engaged by TMS-induced discomfort and deriving neural signatures that generalize across stimulation sites. Second, prior work has largely overlooked potential population differences. Empirical evidence indicates that individuals with depression and anxiety show marked alterations in brain systems involved in pain processing (20, 21). Including both healthy and clinical populations is thus essential to determine whether they show distinct neural signatures of TMS-induced discomfort. Such knowledge would help disentangle direct neuromodulatory effects from symptom-related biases in discomfort processing. Finally, traditional univariate analyses assess neural correlates of a behavioral measure for each brain region independently, which ignores distributed and joint patterns of activity across brain regions and limits sensitivity to capture complex brain-behavior relationships (22). Multivariate approaches, such as the principal component analysis (PCA) combined with the sparse canonical correlation analysis (sCCA) established in our prior work (23–25), provide a powerful computational framework for identifying coordinated neural patterns across multiple regions and accounting for shared variance to isolate the neural patterns most relevant to discomfort (26).

Addressing these gaps, we conducted a large-scale concurrent TMS-fMRI study to map neural responses to TMS-induced discomfort across 11 cortical sites in two cohorts: healthy individuals (N1 = 80) and participants with elevated depression and/or anxiety symptoms (N2 = 85), as assessed by the Patient Health Questionnaire-9 (PHQ-9) or Generalized Anxiety Disorder-7 (GAD-7) scale. Participants completed up to 11 TMS-fMRI runs, each targeting a different cortical site, yielding 1,535 runs in total. Single TMS pulses were delivered while whole-brain responses were recorded, and TMS-induced discomfort was rated immediately after each run using the Subjective Units of Distress Scale (SUDS) (27). Leveraging the PCA-sCCA framework, we identified distributed neural patterns associated with TMS-induced discomfort, revealing regional engagement of sensorimotor, attentional, default mode and limbic networks, as well as subcortical regions, with both shared and group-specific engagement patterns. Moreover, discomfort-related activity accounts for a substantial proportion of TMS-evoked brain responses. Together, these findings provide the first brain-wide mapping of TMS-induced discomfort, establishing a benchmark for interpreting TMS neural effects and highlighting the need to routinely measure and account for TMS-induced discomfort in research and clinical practice.

## Results

### Variability in TMS-induced discomfort across stimulation sites and individuals

A total of 165 participants completed the TMS-fMRI experiment and were included in final analyses for this study (see Methods). They included two groups: the healthy group (PHQ-9 ≤ 5 and GAD-7 ≤ 5; N1 = 80; 47 females; mean age ± S.D. = 31.7 ± 10.6 years) and the symptomatic group with depression and/or anxiety symptoms (PHQ-9 ≥ 10 or GAD-7 ≥ 10; N2 = 85; 57 females; mean age ± S.D. = 31.2 ± 10.6 years) in the clinical range but not confirmed with a formal diagnostic interview. During the experiment, participants received a series of single pulses of TMS inside the MRI scanner interleaved with whole-brain BOLD response recordings (**Fig. 1A**). Each participant completed up to 11 concurrent TMS-fMRI runs, each targeting a different cortical site (**Fig. 1B**). These sites included the left and right frontal pole (L/R FP), left and right anterior middle frontal gyrus (L/R aMFG), left and right posterior middle frontal gyrus (L/R pMFG), right inferior frontal joint (R IFJ), right frontal eye field (R FEF), right pre-supplementary motor area (R preSMA), right primary motor cortex (R M1), and right inferior parietal lobule (R IPL). The coordinates of these stimulation sites were selected from the prior literature (28, 29) and intended for investigating causal circuit engagement within a broad range of cortical networks (**Table 1; Table S1**–**S2**) (see Methods). Immediately after each run, participants were instructed to rate their discomfort to TMS using the SUDS, ranging from 0 (indicating “no discomfort/pain”) to 100 (indicating “the worst pain that you can imagine”).

**Figure 1.**
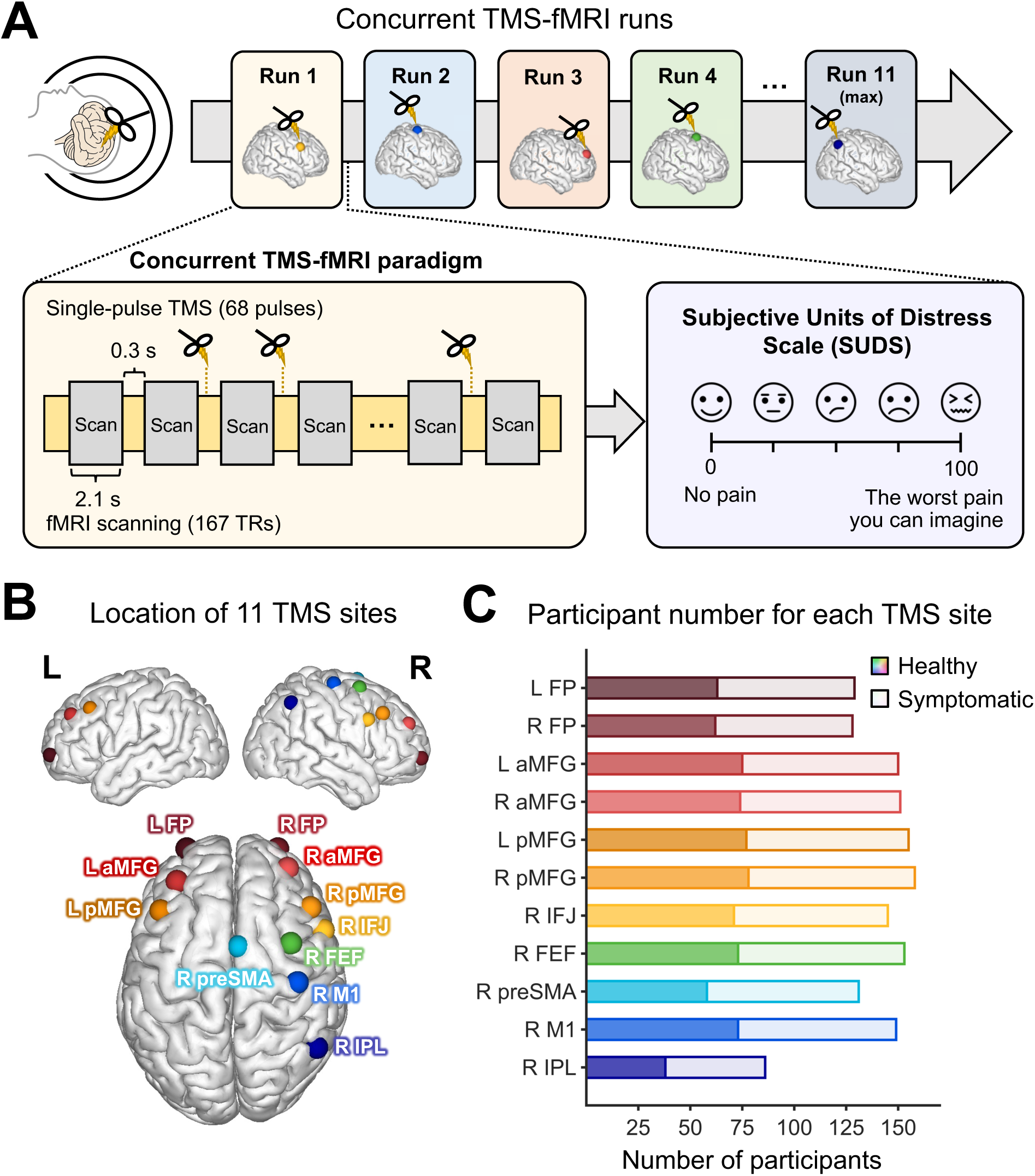
Experimental protocol, stimulation sites, and participant distribution. **(A)** Concurrent TMS-fMRI experimental protocol. Participants completed up to 11 TMS-fMRI runs, with each run targeting a different cortical site with single-pulse TMS inside the MRI scanner interleaved with whole-brain fMRI scans. Immediately after each run, participants orally reported the TMS-induced discomfort using the Subjective Units of Distress Scale (SUDS), with perceived pain or distress from 0 (“no discomfort or pain”) to 100 (“the worst pain that you can imagine”). **(B)** Cortical locations of 11 stimulation targets. Left/right frontal pole (L/R FP), left/right anterior middle frontal gyrus (L/R aMFG), left/right posterior middle frontal gyrus (L/R pMFG), right inferior frontal junction (R IFJ), right frontal eye field (R FEF), right pre-supplementary motor area (R preSMA), right primary motor cortex (R M1), and right inferior parietal lobule (R IPL). The corresponding MNI coordinates of each site are listed in **Table 1**. **(C)** The number of participants in healthy and symptomatic groups completed each TMS-fMRI run.

**Table 1.** Participants’ demographic information and MNI coordinates for each TMS site.

| TMS sites | Number of participants | Healthy group | Symptomatic group | Female | Male | Age (S.D.) | Years of Education (S.D.) | MNI coordinates |  |  |
| --- | --- | --- | --- | --- | --- | --- | --- | --- | --- | --- |
|  |  |  |  |  |  |  |  | x | y | z |
| L FP | 129 | 63 | 66 | 78 | 51 | 31.8(11.1) | 15.9(2.1) | -24 | 60 | -2 |
| R FP | 128 | 62 | 66 | 76 | 52 | 32.1(11.0) | 16.0(2.1) | 24 | 60 | -2 |
| L aMFG | 150 | 75 | 75 | 93 | 57 | 31.0(10.2) | 15.8(2.0) | -32 | 42 | 34 |
| R aMFG | 151 | 74 | 77 | 96 | 55 | 31.4(10.6) | 15.9(2.0) | 30 | 50 | 26 |
| L pMFG | 155 | 77 | 78 | 97 | 58 | 31.6(10.7) | 15.9(2.0) | -38 | 22 | 38 |
| R pMFG | 158 | 78 | 80 | 100 | 58 | 31.4(10.6) | 15.9(2.0) | 46 | 26 | 38 |
| R IFJ | 145 | 71 | 74 | 92 | 53 | 31.6(10.8) | 15.8(2.1) | 46 | 8 | 28 |
| R FEF | 153 | 73 | 80 | 100 | 53 | 31.3(10.6) | 15.8(2.0) | 34 | 6 | 62 |
| R preSMA | 131 | 58 | 73 | 82 | 49 | 30.4(10.1) | 15.8(2.1) | 6 | 2 | 68 |
| R M1 | 149 | 73 | 76 | 95 | 54 | 31.5(10.6) | 15.9(2.0) | 40 | -18 | 64 |
| R IPL | 86 | 38 | 48 | 54 | 32 | 29.6(9.8) | 15.8(2.0) | 48 | -54 | 46 |

A linear mixed-effects model was performed to examine whether the SUDS rating varied across stimulation site, group (healthy vs. symptomatic), and participant demographics (gender and age). Results revealed a statistically significant main effect of site (linear mixed-effects model, *F*_(10, 1290)_ = 9.64, *p* = 1.23 × 10^-15^, partial *η*^2^ = .082). Post-hoc comparisons showed a graded pattern of SUDS ratings across sites, decreasing from frontal to posterior and from inferior to superior regions: SUDS ratings were highest at L/R FP, followed by L/R aMFG, L/R pMFG, and R IFJ; then R FEF; and lowest at R preSMA, R M1, R IPL (*p*_FDR_ < .001) (**Fig. 2A/B**). Consistent results were observed when the cross-site comparison was conducted separately for the healthy and symptomatic groups (**Fig. S2**). Additionally, significant differences were observed in the MFG sites between left vs. right hemispheres, with higher SUDS ratings at R aMFG/pMFG than at L aMFG/pMFG (*p*_FDR_ < .011). No other main effects or interactions were significant (*p*s > .16). In particular, no significant differences were observed in the SUDS ratings between the two groups at any stimulation site (*p*s > .15).

**Figure 2.**
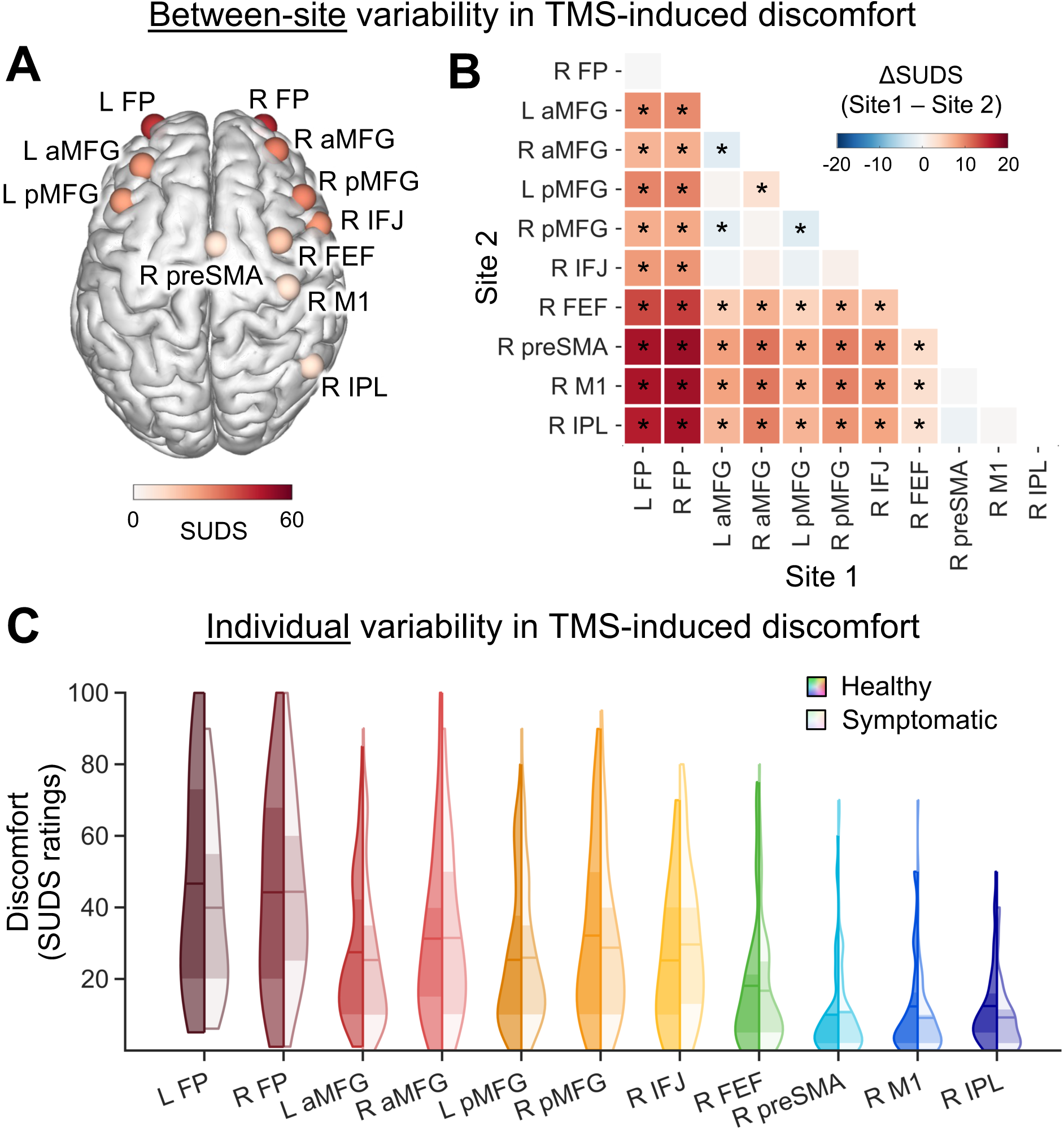
Variability in TMS-induced discomfort across stimulation sites and individuals. **(A**, **B)** Discomfort variability across stimulation sites. **A,** group-averaged discomfort rating at each stimulation site. **B,** pairwise differences in discomfort between sites, with statistically significant between-site differences marked by asterisks (*p*_FDR_ < 0.05). Discomfort is highest at L/R FP, followed by L/R aMFG, L/R pMFG, and R IFJ; then R FEF; and lowest at R preSMA, R M1, R IPL. Additionally, discomfort is higher at R aMFG/pMFG than at L aMFG/pMFG. Here, SUDS ratings from the two groups were pooled. Similar results were observed when each group were analyzed separately (**Fig. S2**). **(C)** Substantial individual variability in discomfort within each group for each stimulation site. Mean values are marked by the horizontal bars. The interquartile range (25^th^ – 75^th^ percentile) is shown by the darker shading. Abbreviations: SUDS, Subjective Units of Distress Scale; FP, frontal pole; aMFG, anterior middle frontal gyrus; pMFG, posterior middle frontal gyrus; IFJ, inferior frontal junction; FEF, frontal eye field; preSMA, pre-supplementary motor area; M1, primary motor cortex; IPL, inferior parietal lobule.

The distribution of SUDS ratings of all participants for each stimulation site and each group is presented in **Fig. 2C**. Substantial interindividual variability was observed within each site (**Fig. 2C, Table S3**). For example, at the R FP, which showed the highest group-average SUDS (mean = 44.3) across all sites, participant’s ratings spanned a wide range, with the lower quartile (25^th^ percentile) reporting SUDS scores ≤ 20, whereas the upper quartile (75^th^ percentile) reporting scores ≥ 65. Complete statistical details for each site and group, including the minimum, maximum, interquartile range, mean and standard deviation, are provided in **Table S3**. Together, the variability observed across sites and individuals provides the wide range of discomfort necessary for modeling the neural correlates of TMS-induced discomfort.

### Selection of a computational framework for modeling TMS-induced discomfort

To identify the computational framework that best captures neural signatures of TMS-induced discomfort, we compared the performance of several multivariate approaches relating the TMS-evoked brain responses to SUDS ratings. Analyses were performed separately for each group using all available TMS-fMRI runs across participants (healthy group: n1 = 741 runs; symptomatic group: n2 = 793 runs). For each run, TMS-evoked brain responses were extracted from 246 cortical and subcortical ROIs defined by the Brainnetome (BN) atlas (30), yielding one high-dimensional multivariate brain-response sample per run for subsequent modeling (**Fig. 3A**).

**Figure 3.**
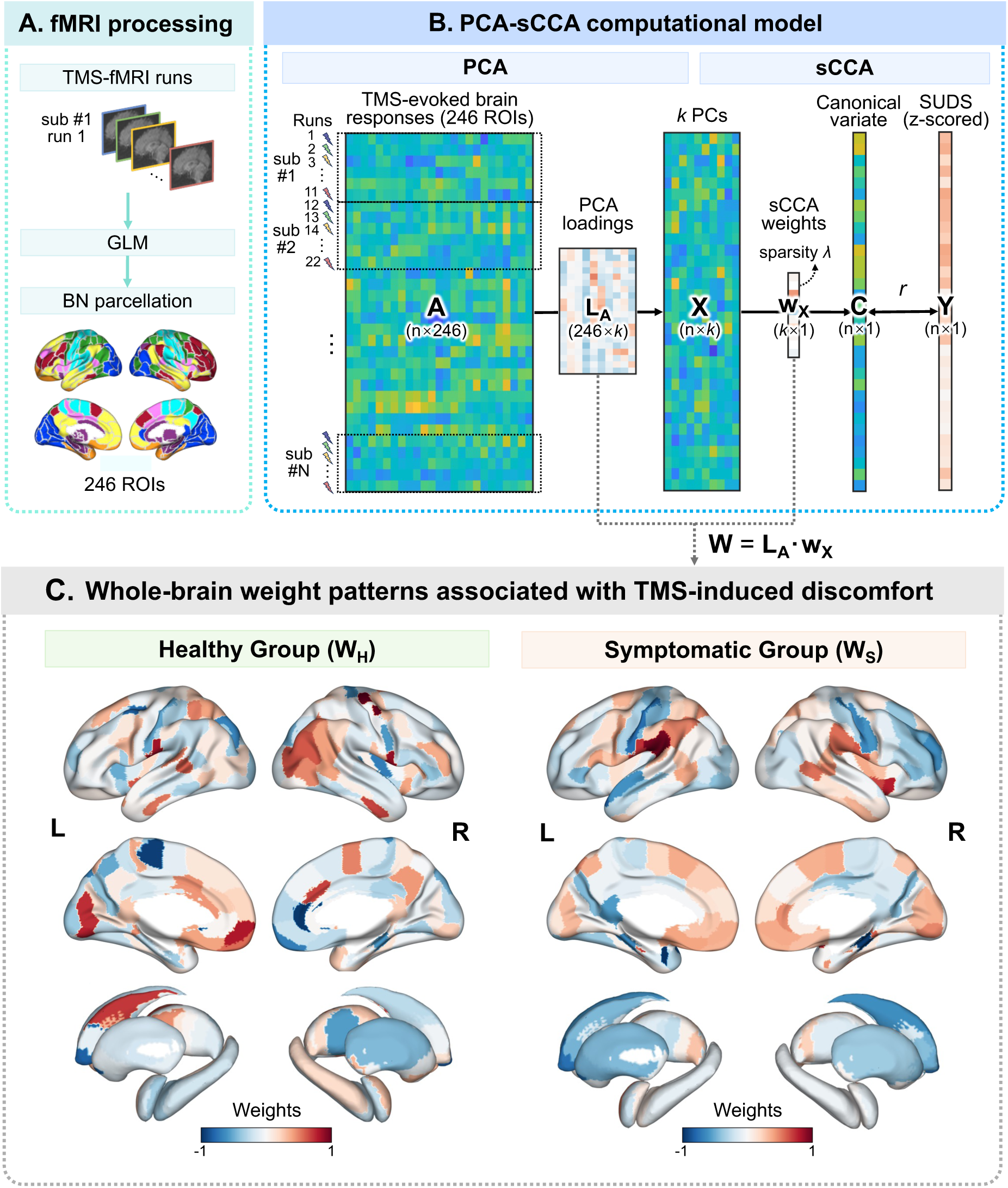
Framework for identifying neural signatures of TMS-induced discomfort. (A) fMRI processing. Each TMS-fMRI run for each participant was first preprocessed and TMS-evoked brain responses were estimated with a generalized linear model (GLM) and extracted with the Brainnetome (BN) atlas that consists of 246 ROIs. These samples of brain responses together with SUDS (healthy group: N1 = 80 subjects, n1 = 741 runs; symptomatic group: N2 = 85 subjects, n2 = 793 runs) were entered into the PCA-sCCA model. (B) PCA-sCCA computational model. Principal component analysis (PCA) was first used to reduce dimensionality and capture dominant variance in TMS-evoked brain responses (𝑨; n × 246), yielding *k* retained principal components (PCs, 𝑿; n × *k*) through a 246 × *k* PCA loading matrix (𝑳_𝑨_). Sparse canonical correlation analysis (sCCA) was then applied to the *k* PCs to identify a linear weighted combination of PCs, defined by the sCCA weight vector 𝒘_𝑿_ (*k* × 1), yielding a canonical variate for brain responses (𝑪 = 𝑿 · 𝒘_𝑿_; n × 1) that maximize the correlation (*r*) with SUDS ratings (𝒀; n × 1). The sparsity of the sCCA weight vector was defined by *λ*, controlling the number of nonzero weights in the sCCA solution. The correlation *r* quantified the strength of the brain-discomfort association. Model performance was evaluated using ten rounds of ten-fold cross-validation (see **Fig. S3A**). The model was performed separately for the healthy and symptomatic groups. (C) Whole-brain weight patterns representing the brain-discomfort association in the healthy group (left) and the symptomatic group (right). These whole-brain patterns (healthy group: 𝑾_𝑯_; symptomatic group: 𝑾_𝑺_) were derived as the product of the PCA loading matrix (𝑳_𝑨_) and the sCCA weight vector (𝒘_𝑿_) for each group. The hyperparameters were determined for each group as the combination yielding the highest cross-validated model generalizability (see Results and Methods; **Fig. S3B–C**).

We first evaluated a PCA-sCCA framework established in our prior work (23–25). In this framework (**Fig. 3B**), PCA was first applied to the high-dimensional multivariate TMS-evoked brain-response data (𝑨; n × 246) to estimate PCA loadings (𝑳_𝑨_ ; 246 × *k*) and derive low-dimensional principal components (PCs) capturing the dominant variance in brain responses (𝑿; n × *k*). The number of retained PCs (*k*) was treated as a tunable hyperparameter. sCCA was then applied to the retained PCs to identify multivariate associations between TMS-evoked brain responses and TMS-induced discomfort. This analysis constructed a weighted combination across PCs, defining a single canonical variate of brain responses (𝑪 = 𝑿 · 𝒘_𝑿_; n × 1) that was maximally correlated with SUDS ratings (𝒀; n × 1). The resulting sCCA weights (𝒘_𝑿_; *k* × 1) specifies how strongly each retained PC in 𝑿 contributes to the brain canonical variate that best relates to discomfort. The sparsity parameter (*λ*), controlling the number of nonzero weights in the sCCA solution to limit model complexity and improve model interpretability and generalizability, was also treated as a tunable hyperparameter. To optimize model hyperparameters and to evaluate the model performance, ten rounds of ten-fold cross-validation (CV) were performed (see Methods). Within each CV fold, 90% of participants were assigned to the training set and 10% to the held-out validation set (**Fig. S3A**). The PCA loadings and sCCA weights estimated from the training data were subsequently applied to the validation data to generate a cross-validated brain canonical variate. The correlation (*r*_CV_) between the cross-validated brain variate and SUDS ratings served as a cross-validated estimate of model generalizability.

We found that, in the healthy group, retaining *k* = 160 PCs in the PCA and setting *λ* = 0.25 in the sCCA (**Fig. S3B**) provided optimal hyperparameters, achieving the highest and significant cross-validated generalizability across CVs (*r*_CV_ = 0.28 ± 0.02, mean ± S.D., *p*_permutation_ < .001) (**Fig. S4A–B**). With these hyperparameters, the overall PCA-sCCA weight pattern, defined as the product of the PCA loadings (246 × *k*) and sCCA weights (*k* × 1), showed high stability across CV folds and rounds, as quantified by intra-class correlation coefficient (ICC = 0.83 ± 0.03, mean ± S.D.) (see Methods). In the symptomatic group, the optimal hyperparameters were *k* = 50 PCs and *λ* = 0.20 (**Fig. S3C**), yielding similarly robust generalizability (*r*_CV_ = 0.25 ± 0.01, mean ± S.D., *p*_permutation_ < .001) (**Fig. S4C–D**) and high stability (ICC = 0.90 ± 0.03).

To further assess the comparative performance of the PCA-sCCA framework, we evaluated several common multivariate computational frameworks, including sCCA alone, sparse partial least squares (sPLS) alone, and the PCA-sPLS combination. Using the same cross-validation strategy, PCA-sCCA consistently outperformed these alternatives in both cross-validated generalizability (*r*_CV_) and a combined metric of generalizability (*r*_CV_) and stability (ICC) (31) (**Fig. S5; Table S4**). These results support the selection of PCA-sCCA as the primary framework for modeling brain-discomfort associations.

### Neural signatures associated with TMS-induced discomfort

Given the superior performance of the PCA-sCCA framework, we next applied it to the full samples to characterize neural signatures of TMS-induced discomfort. Analyses were conducted separately for each group using the optimal hyperparameters identified above (healthy group: *k* = 160 PCs, *λ* = 0.25; symptomatic group: *k* = 50 PCs, *λ* = 0.20; **Fig. S3B–C**). Building on the PCA-sCCA framework described above, we computed whole-brain weight patterns representing the association between TMS-evoked brain responses and TMS-induced discomfort as the product of the PCA loadings and the sCCA weights, thereby projecting the brain-discomfort association identified in the reduced PCA space back into the original 246-ROI space. Each element represents the contribution of a given ROI to the overall brain-discomfort association in the whole-brain space (**Fig. 3C**). Weight magnitude indicates the strength of the ROI’s contribution, whereas the sign reflects its direction, with positive weights indicating ROIs whose brain responses increase with higher SUDS ratings and negative weights indicating the opposite pattern. To identify the critical regions that contributed most strongly to the brain-discomfort association, we focused on the top 15 ROIs (∼6% of all regions) ranked by absolute weight magnitude within each group (**Fig. 4**), following our prior studies (25, 32, 33) . Permutation test (10,000 iterations), conducted separately for each group, demonstrated that all these top ROIs for two groups exhibited statistically significant weights (*p*_permutation_ < 0.05, FDR corrected across ROIs) or marginally significant (*p*_permutation_ < 0.07, FDR corrected) weights (**Fig. S6; Table S5–S6**). The spatial distribution and network assignment of these top ROIs were then examined, followed by between-group comparisons of ROI-wise weights.

**Figure 4.**
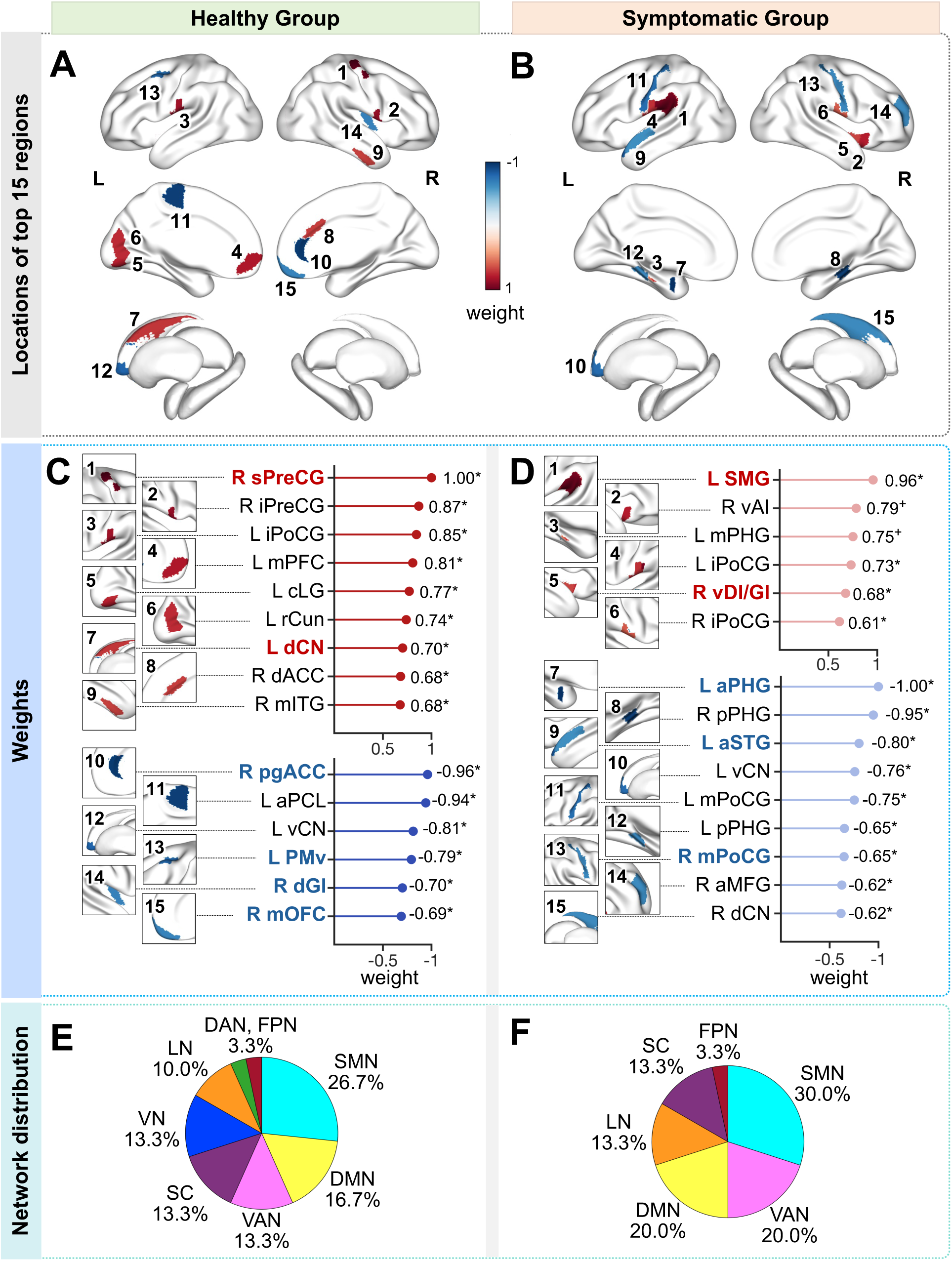
Locations, weights and network distribution of top 15 ROIs most strongly associated with TMS-induced discomfort in the healthy and symptomatic groups. **(A, B)** Spatial locations of the top 15 ROIs with the largest absolute weights identified by the PCA-sCCA model in the healthy group (**A**) and the symptomatic group (**B**). (**C**, **D**) Weights of top 15 ROIs for each group. ROIs with positive and negative weights are listed separately. In the healthy group (**C**), all ROI weights were statistically significant by permutation test (10,000 iterations, *p*_permutation_ < 0.05, FDR corrected; indicated by asterisks) (**Fig. S6A, Table S5**). These weights were then compared with the corresponding weights in the symptomatic group. Six ROIs exhibited significant cross-group differences by permutation test (*p*_permutation_ < 0.05, FDR corrected; **Table S7**), indicated by colored text. In the symptomatic group, all ROI weights were either statistically significant (*p*_permutation_ < 0.05, FDR corrected; indicated by asterisks) or marginally significant (*p*_permutation_ < 0.07, FDR corrected; indicated by crosses) (**Fig. S6B, Table S6**). Five ROIs showed significant (*p*_permutation_ < 0.05, FDR corrected) or marginal significant (*p*_permutation_ < 0.07, FDR corrected) cross-group differences (**Table S8**), indicated by colored text. (**E**, **F**) Network distribution of top 15 ROIs for each group. Each ROI were first assigned to one or more of the seven functional networks (**Table S5/S6;** see Methods), and proportion of top ROIs assigned to each network was then quantified. In the healthy group (**E**), the top ROIs were primarily distributed across the SMN and DMN, followed by the VAN, SC, VN and LN. In the symptomatic group (**F**), the top ROIs were primarily distributed across the SMN, VAN, and DMN, followed by the LN and SC. Abbreviations: ROIs: aMFG = anterior middle frontal gyrus; aPCL = anterior paracentral lobule; aPHG = anterior parahippocampal gyrus; aSTG = anterior superior temporal gyrus; cLG = caudal lingual gyrus; dACC = dorsal anterior cingulate cortex; dCN = dorsal caudate nucleus; dGI = dorsal granular insula; iPoCG = inferior postcentral gyrus (parietal operculum, PO; secondary somatosensory cortex, S2); iPreCG = inferior precentral gyrus; mITG = middle inferior temporal gyrus; mOFC = medial orbitofrontal cortex; mPFC = medial prefrontal cortex; mPHG = middle parahippocampal gyrus; mPoCG = middle postcentral gyrus (primary somatosensory cortex, S1); pgACC = pregenual anterior cingulate cortex; PMv = ventral premotor cortex; pPHG = posterior parahippocampal gyrus; rCun = rostral cuneus; SMG = supramarginal gyrus; sPreCG = superior precentral gyrus; vAI = ventral anterior insula; vCN = ventral caudate nucleus; vDI/GI = ventral dysgranular/granular insula. Networks: DAN = dorsal attention network; DMN = default mode network; FPN = frontoparietal control network; LN = limbic network; SMN = somatomotor network; VAN = ventral attention network; VN = visual network; SC = subcortical regions.

These top ROIs were distributed across cortical and subcortical regions (**Fig. 4**). In the healthy group (**Fig. 4A/C**), they included somatomotor and premotor areas, including the left postcentral gyrus (PoCG), right precentral gyrus (PreCG) and left anterior paracentral lobule (aPCL), left ventral premotor cortex (PMv); occipital areas, including the left caudal lingual gyrus (cLG), left rostral cuneus gyrus (rCun); medial frontal and limbic regions, including the left medial prefrontal cortex (mPFC), right medial orbitofrontal cortex (mOFC), right dorsal and pregenual anterior cingulate cortex (dACC, pgACC); and the right dorsal insula, right middle inferior temporal gyrus (mITG), and left caudate nucleus (CN). In the symptomatic group (**Fig. 4B/D**), the top ROIs also included somatomotor areas, including the bilateral PoCG and left supramarginal gyrus (SMG); temporal regions, including the left anterior superior temporal gyrus (aSTG), bilateral parahippocampal gyrus (PHG); as well as the right ventral insula, left aMFG, and bilateral CN.

These top ROIs spanned multiple functional networks, including the somatomotor, attention, default mode, limbic networks, as well as subcortical regions (**Table S5–S6**). We next quantified the network distribution among these top ROIs separately for each group. In the healthy group, the top ROIs were primarily distributed across the somatomotor (26.7%) and default mode (16.7%) networks, followed by the ventral attention network, subcortical regions, and visual network, each accounting for 13.3% of top ROIs, and the limbic network (10.0%). In the symptomatic group, the top ROIs were also primarily distributed across the somatomotor (30.0%), default mode (20.0%) and ventral attention (20.0%) networks, followed by the limbic network and subcortical regions, each accounting for 13.3% of top ROIs.

Among the top ROIs, two were engaged in TMS-induced discomfort in both healthy and symptomatic groups, including the left inferior PoCG and left ventral CN, whereas the remaining 13 ROIs differed between groups. Between-group comparisons further revealed group-specific regional engagement, reflected in significant differences in ROI weights. Specifically, six showed statistically greater weights in the healthy group, including the left PMv, right mOFC, right superior PreCG, right dorsal granular insula, right dACC, and left dorsal CN (*p*_permutation_ < 0.05, FDR corrected; **Fig. 4C, Table S7**). In contrast, five ROIs showed significantly or marginally greater weights in the symptomatic group, including the left aSTG and left SMG (*p*_permutation_ < 0.05, FDR corrected), as well as the left anterior PHG, right middle PoCG, and right ventral dysgranular/granular insula (*p*_permutation_ < 0.07, FDR corrected; **Fig. 4D, Table S8**). No significant between-group differences were observed for the remaining ROIs (*p*_permutation_ > 0.10, FDR corrected).

### Proportion of TMS-evoked brain responses attributable to discomfort-related signatures

To quantify the magnitude of discomfort-related influence in TMS-evoked brain responses, we projected each sample’s ROI-wise TMS-evoked brain responses onto the whole-brain discomfort-related weight map to obtain the discomfort-related response component and estimated the average of the proportion it accounted for in the TMS-evoked brain responses across samples (**Fig. 5A**; see Methods). This proportion was computed for each ROI within each group separately. In the healthy group, the discomfort-related responses accounted for 12.3% ± 2.4% (mean ± S.D.) of the TMS-evoked responses across the top 15 ROIs, and 3.7% ± 2.8% (mean ± S.D.) across the remaining ROIs (**Fig. 5B**). In the symptomatic group, the discomfort-related responses accounted for 25.2% ± 4.9% (mean ± S.D.) across the top 15 ROIs and 8.2% ± 5.7% (mean ± S.D.) across the remaining ROIs (**Fig. 5B**).

**Figure 5.**
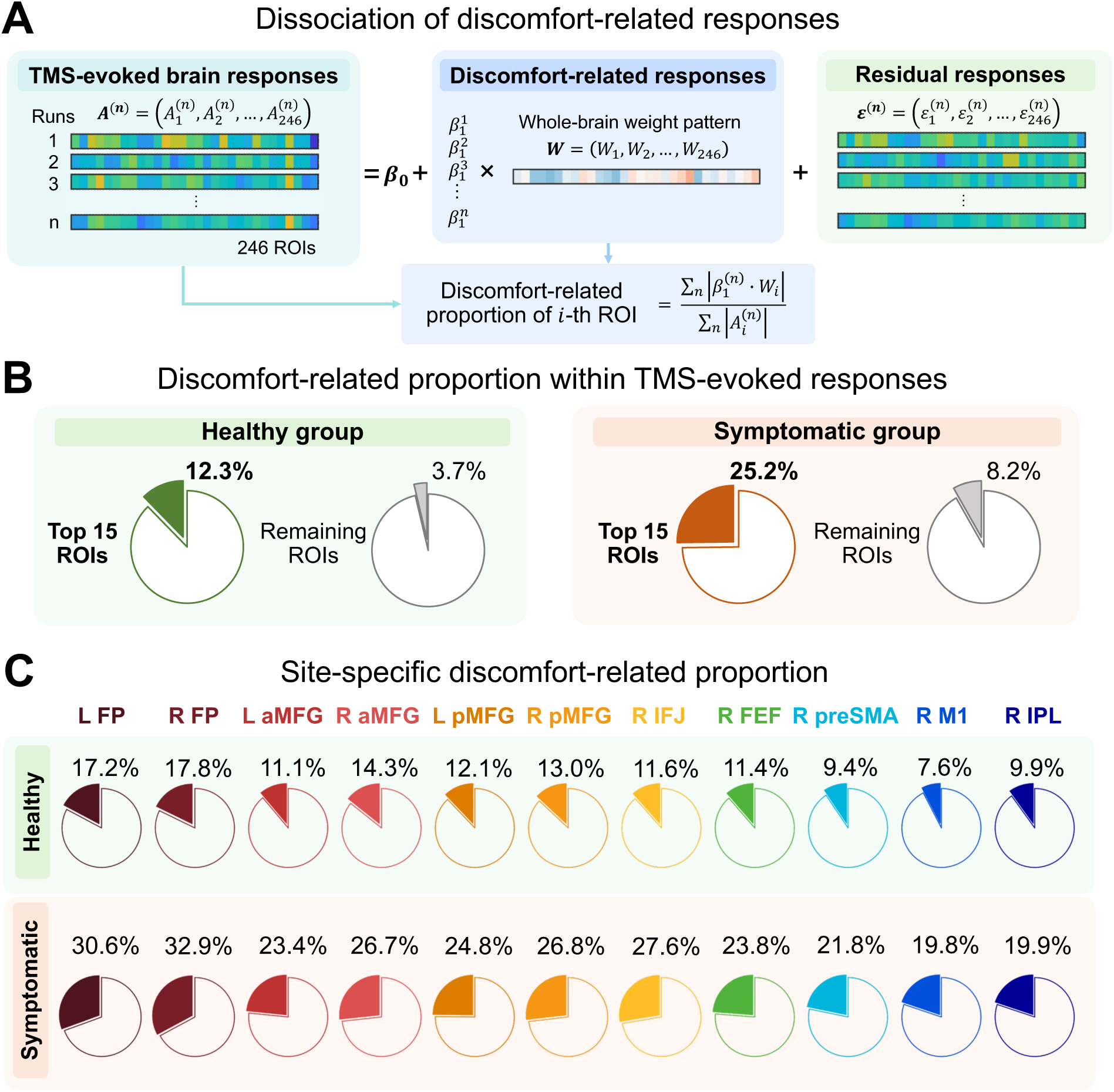
Discomfort-related proportion of TMS-evoked brain responses in the healthy and symptomatic groups. **(A)** Computation of discomfort-related proportion. For each sample, TMS-evoked brain responses 𝑨^(&)^were projected onto the whole-brain discomfort-related weight vector 𝑾, yielding a discomfort-related component 𝛽^(&)^ · 𝑾. The discomfort-related proportion was quantified as the relative magnitude of this discomfort-related component to the original TMS-evoked brain response at each ROI and then averaged across samples. **(B)** Discomfort-related proportion in the healthy and symptomatic groups. In the healthy group, discomfort-related responses accounted for an average of 12.3% of TMS-evoked brain responses across the top 15 ROIs with the largest weight magnitudes, and 3.7% across the remaining ROIs. In the symptomatic group, discomfort-related responses accounted for an average of 25.2% of TMS-evoked brain responses across the top 15 ROIs, and 8.2% across the remaining ROIs. **(C)** Site-specific discomfort-related proportion within the top 15 ROIs. In both groups, the proportion of discomfort-related activity showed a graded pattern across stimulation sites, decreasing from frontal to posterior and from inferior to superior sites. The symptomatic group exhibited higher proportions than the healthy group at all TMS sites.

Given substantial differences in TMS-induced discomfort across stimulation sites, we further evaluated the magnitude of discomfort-related influence for each TMS site separately. Consistent with the pattern observed in SUDS ratings, the proportion of discomfort-related activity showed a graded pattern across stimulation sites, decreasing from frontal to posterior and from inferior to superior sites, in both groups (**Fig. 5C; Table S9**). Specifically, the L/R FP showed the highest proportions (Healthy group: L FP, 17.2% ± 3.9%, mean ± S.D., R FP, 17.8% ± 3.8%; Symptomatic group: L FP, 30.6% ± 6.9%, R FP, 32.9% ± 7.0%), whereas R M1 showed the lowest proportions (Healthy group: R M1, 7.6% ± 1.5%, mean ± S.D.; Symptomatic group: R M1, 19.8% ± 3.3%). Same as the overall pattern, there is stronger discomfort-related influence in the symptomatic group than the healthy group at all TMS sites (**Fig. 5C; Table S9**).

### Group specificity of identified discomfort-related neural signatures

To test whether the identified neural signatures were specific to each group, we conducted two complementary analyses. First, we removed the within-group discomfort-related component response from TMS-evoked brain responses and re-evaluated the correlation between the new brain canonical variate derived from residual TMS-evoked brain responses and SUDS ratings. We expected that if the identified neural signatures robustly accounted for the observed brain-discomfort association in each group, removing it would significantly reduce this correlation. Consistent with this expectation, the original significant correlations between the brain canonical variate and SUDS ratings in both the healthy group (*r* = 0.65, *p*_permutation_ < .001) and the symptomatic group (*r* = 0.43, *p*_permutation_ < .001), based on permutation testing (10,000 iterations), were markedly reduced after removal (healthy group: *r* = 0.43, Δ*r* = -0.21; *z*-test: *z* = 6.06, *p* = 6.68 × 10^-10^; symptomatic group: r = 0.26, Δ*r* = -0.17; *z* = 3.85, *p* = 5.87 × 10^-5^) (**Fig. S7**), and were no longer significant (healthy: *p*_permutation_ = .90; symptomatic: *p*_permutation_ = .36).

Second, we removed each group’s discomfort-related component response from the other group’s TMS-evoked brain responses. We expected that if the identified neural signatures were group-specific, cross-group removal would not eliminate the observed brain-discomfort association. As predicted, significant correlations remained robust in both the healthy group (*r* = 0.63, *p*_permutation_ < .001) and the symptomatic group (*r* = 0.43, *p*_permutation_ < .001), with only minimal reductions relative to original correlations (healthy group: Δ*r* = -0.02, *z*-test: *z* = 0.60, *p* = .28; symptomatic group: Δ*r* < 0.01; *z*-test: *z* = 0.01, *p* = .50) (**Fig. S7**). Together, these results indicate that the identified discomfort-related neural signatures are largely group-specific rather than interchangeable across populations.

## Discussion

In this study, we aimed to characterize the whole-brain neural correlates of TMS-induced discomfort using a large concurrent TMS-fMRI dataset comprising 165 participants and 11 cortical stimulation sites. Leveraging a PCA-sCCA computational framework, we demonstrated that brain responses from distributed neural patterns were associated with subjective ratings of TMS-induced discomfort. These patterns broadly engage cortical regions spanning sensorimotor, attentional, limbic and default mode networks, as well as subcortical regions, with both shared and population-specific regional engagement. Moreover, a substantial proportion of TMS-evoked neural responses was attributable to discomfort-related activity, accounting for approximately 12% and 25% of TMS-evoked responses in the healthy and elevated-symptom groups, respectively. This proportion also varied across stimulation sites, with higher proportions observed at more frontal and inferior sites. Together, these findings provide the first brain-wide mapping of TMS-induced discomfort, highlighting discomfort-related processing as a non-negligible confounding factor in interpreting neuromodulatory effects of TMS. Importantly, the identified patterns offer a practical benchmark for examining, accounting for, and regressing out discomfort-related responses in observed TMS-evoked brain responses. We therefore call for more attention to the routine assessment, reporting, and control of TMS-induced discomfort to improve the interpretability, reproducibility, and translational validity of TMS applications in both basic research and clinical practice.

A major finding of this study is that TMS-induced discomfort was reflected in TMS-evoked responses across a distributed set of cortical and subcortical regions implicated in sensory, nociceptive, attentional, and affective processes. These regions included areas within the somatomotor network, such as the primary and secondary somatosensory areas (S1, S2), posterior insula, and motor and premotor areas, which have been implicated in the sensory processing of pain or aversive stimuli (34, 35). Specifically, S1 and S2 are well established in mapping head-related sensations (36, 37) and might provide somatotopically appropriate encoding of peripheral scalp sensations evoked by TMS pulses, whereas the posterior insula serves as a major cortical target of spinothalamic projections conveying nociceptive signals (36, 37). The engagement of motor and premotor regions likely reflects sensory-motor coupling, which facilitates sensory processing and/or preparatory motor responses to aversive stimuli (38, 39). The discomfort-related signature also extended to occipital regions. Because no visual stimulus was presented in our experiment, activities in the visual regions is unlikely to reflect the visual input. Instead, it may reflect multisensory integration processes involving visual, somatosensory, and vestibular systems (40) during the processing of uncomfortable bodily sensations evoked by TMS pulses.

Discomfort-related neural signatures further engaged higher-order cortical regions spanning the ventral attention, default mode and limbic networks, as well as subcortical regions. As expected, the anterior insula and dACC, two key nodes of the ventral attention network, were engaged. Prior work has implicated the anterior insula in the detection of salient bodily sensations (38, 41), and the dACC in pain processing (42) and the integration of sensory and nociceptive signals into conscious awareness (38, 41). Their engagement here therefore likely reflects the detection and perceptual processing of pain sensations elicited by TMS over the scalp. The engagement of regions within the default mode and limbic networks may further support the appraisal and contextual interpretation of these sensations. For example, PHG, a representative region in the default mode network, may support the integration of current discomfort with contextual information or prior aversive memories (43, 44). In addition, medial prefrontal and limbic regions, including the mPFC, mOFC and pgACC, have been implicated in supporting affective and cognitive appraisal of discomfort (45). Beyond these, discomfort-related signatures involved cortical regions within the fronto-parietal control network (e.g., aMFG), which may reflect anticipatory and/or regulatory processes related to discomfort, potentially via cognitive control and emotional regulation (46, 47). In addition, subcortical regions, particularly the CN, were also involved and may contribute to nociceptive modulation (48) and relief of TMS-induced discomfort. Taken together, TMS-induced discomfort engages distributed brain regions that jointly support a multidimensional process encompassing perception, evaluation, and modulation of discomfort (34, 35). Notably, the organization of these distributed patterns aligns with well-established neural mechanisms of acute pain processing observed in classical laboratory paradigms using mechanical pressure, thermal stimuli, or intraepidermal electrical stimulation (45, 49, 50). This correspondence suggests that TMS-induced discomfort is processed in a manner analogous to nociceptive stimuli experienced as pain.

Brain patterns associated with TMS-induced discomfort showed both shared and population-specific regional engagement across healthy individuals and participants with negative affect symptoms. Aside from the shared engagement of the left inferior PoCG and left ventral CN, both of which have been widely implicated in pain and discomfort processing (13), the two groups showed substantial differences in the broader regional engagement. Within the somatomotor network, symptomatic participants showed weaker engagement of motor regions, i.e., PreCG, but broader and stronger engagement of somatosensory regions, i.e., bilateral PoCG. In higher-order cortical regions, symptomatic participants showed more engagement of attentional regions, particularly the right anterior insula, and of memory-associated regions, including the PHG and aSTG. This pattern may indicate increased attentional allocation toward aversive stimulation (51) and a greater tendency to associate current discomfort with prior negative experiences or form new unpleasant memories (43, 44). Meanwhile, symptomatic participants showed less engagement of frontal and limbic regions, suggesting altered appraisal of discomfort-related signals. Consistent with these findings, prior neuroimaging studies have reported structural and functional abnormalities within these regions in depression (52) and anxiety (53). In subcortical regions, the symptomatic participants showed less engagement of the left dorsal CN, suggesting altered modulatory processing of discomfort (48). Together, these findings suggest differential engagement of distributed perception-evaluation-modulation systems during discomfort processing across populations (34, 39). Despite these neural engagement differences between groups, interestingly, we did not observe significant group differences on subjective ratings of TMS-induced discomfort. This finding is consistent with prior meta-analytic behavioral evidence showing no systematic group differences in pain perception between healthy individuals and those with depression (54). Our behavioral and neural results together suggest that different populations process discomfort differently despite similar subjective peripheral sensations.

By quantifying a substantial extent of discomfort-related responses observed in TMS-evoked brain responses, our findings underscore that TMS-induced discomfort constitutes a nontrivial confounding factor when interpreting TMS-evoked neural outcomes. This confounding influence was especially evident for prefrontal stimulation, which elicited stronger discomfort than stimulation of posterior regions, consistent with prior reports, and correspondingly showed a higher proportion of the TMS-evoked brain responses attributable to discomfort-related processing. Given that prefrontal TMS is widely used in both basic and clinical research, particularly in studies of cognitive control (55), emotional regulation (56), and psychiatric disorders (57), these findings indicate that particular caution is warranted when interpreting TMS-evoked neural effects following prefrontal stimulation. This concern is amplified by the substantial overlap between brain regions or circuits commonly targeted or interpreted in prefrontal TMS studies and the distributed discomfort-related neural signatures identified here. For example, several concurrent TMS-fMRI (58, 59) and TMS-intracranial electroencephalography studies (60) have reported ACC or insula activation following prefrontal stimulation and interpreted these responses as evidence of causal engagement via propagation of stimulation-specific neuromodulatory effects. However, in light of our findings, responses in ACC or insula may partly reflect discomfort-related processing rather than exclusively indexing neuromodulatory effects.

Given the population-specific nature of discomfort-related neural patterns identified in this study, caution is further warranted when interpreting between-population differences in TMS-evoked responses (61, 62), particularly when such differences were observed in regions differentially involved in discomfort-related processing across populations. Notably, compared with healthy participants, a higher proportion of TMS-evoked brain responses was attributable to discomfort-related processing in individuals with negative affect symptoms. This difference may arise from heightened attentional engagement and greater allocation of cognitive resources toward processing aversive sensations in symptomatic individuals (51), as suggested by the broader engagement of attentional and memory-related regions associated with TMS-induced discomfort. Accordingly, TMS-evoked brain responses in symptomatic populations warrant careful interpretation. Taken together, our findings indicate that explicitly modeling and controlling for discomfort-related neural responses is critical for accurate interpretation of TMS-evoked neural effects. We therefore emphasize the importance of routinely assessing and controlling discomfort in TMS experiments. When discomfort levels are high, researchers should explicitly incorporate discomfort measures into analytic models, such as treating them as covariates or regressing out discomfort-related responses, to disentangle the discomfort-driven influences from stimulation-specific neuromodulatory effects of interest. The population-specific neural patterns identified here may provide a useful reference for removing discomfort-related confounds, as supported by the attenuation of brain-discomfort associations after regressing out the corresponding pattern.

Future studies could extend the present findings in several ways. First, although our dataset is large, replication across independent research sites using different scanners, TMS coils, and stimulation machines will be essential for establishing the robustness and generalizability of the identified neural patterns. Second, future work should determine whether the discomfort-related neural pattern evoked by the single-pulse stimulation generalizes across other widely used TMS protocols, including repetitive TMS. These repetitive TMS protocols differ in stimulation sequence, duration, and induced sensory experience, and may therefore differentially recruit brain regions or circuits involved in discomfort-related processing. Third, the current study included only participants with depression and/or anxiety symptoms as the clinical group. Given the broad clinical application of TMS in psychiatric, movement, and cognitive disorders (63) and the likelihood that discomfort is processed differently across diagnostic conditions, it is necessary to examine additional clinical populations, such as substance addiction, obsessive-compulsive disorder, dementia, etc. Finally, TMS-EEG and TMS-iEEG studies can adopt the PCA-sCCA framework to identify spatiotemporal signatures of discomfort, thereby providing valuable insights into the rapid temporal dynamics of TMS-induced discomfort processing that are inaccessible with TMS-fMRI.

In conclusion, this study provides the first large-scale characterization of how TMS-induced discomfort is represented in the human brain. Our findings demonstrate that discomfort processing engages distributed, brain-wide neural patterns in a population-specific manner, and further reveal that a substantial fraction of TMS-evoked brain responses might reflect discomfort-related processing. Accordingly, we recommend that future TMS studies routinely assess discomfort induced by TMS and explicitly account for it in analyses, either by regressing out discomfort-related neural patterns or by modeling discomfort ratings as covariates, particularly when discomfort is pronounced. Implementing these practices will improve the interpretability and methodological rigor of TMS research in both clinical applications and basic neuroscience.

## Methods

### Participants

Participants were recruited via advertisements for participation in a study investigating causal connectome mechanisms of the human brain. They were invited to complete 2 neuroimaging visits. Before the visits, an online screening and intake procedure was conducted. Trained doctoral-level clinicians administered clinical interviews to verify the absence of current and past neurological disorders, greater than mild traumatic brain injury (> 30 min loss of consciousness or > 24 hrs post-trauma amnesia), claustrophobia, or regular use of psychiatric medications. To determine clinical grouping, participants completed a comprehensive evaluation, consisting of structured interviews, neuropsychological assessments, and self-report questionnaires. The initial questionnaire collected demographic information, including but not limited to age, sex assigned at birth, ethnicity, and race. Then, several self-report instruments were used to evaluate psychological constructs, including the Patient Health Questionnaire-9 (PHQ-9) for depression screening and the Generalized Anxiety Disorder-7 (GAD-7) for anxiety screening. Participants with both scores ≤ 5 or with either score ≥ 10 were invited for subsequent visits.

During the two visits, participants underwent MRI scanning on two separate days. On Day 1, high-resolution structural images (T1-weighted), resting-state fMRI data, and several task-based fMRI data were acquired. The resting-state and task fMRI data were not used in the present study and are therefore not described further. On Day 2, participants completed up to 11 concurrent TMS-fMRI runs across different stimulation sites. The number of stimulation sites varied across participants due to reasons including 1) overall scan time constraints to complete all the TMS sites, 2) discomfort at certain stimulation sites, 3) spatial constraints within the MRI head coil for participants with bigger head sizes, and 4) the addition of the parietal stimulation site at a later stage of the study. This resulted in different numbers of participants across stimulation sites (**Fig. 1C)**.

The final dataset included a total of 165 adult right-handed participants (104 females, 61 males; mean age = 31.4 ± 10.6 years, S.D.), yielding 1,535 TMS-fMRI runs in total. Based on PHQ-9 and GAD-7 scores, they were categorized into a healthy group (N = 80; 47 females, 33 males; mean age = 31.7 ± 10.6 years, S.D.; PHQ-9 ≤ 5 and GAD-7 ≤ 5) and a symptomatic group experiencing clinically relevant negative affect symptoms (N = 85; 57 females, 28 males; mean age = 31.2 ± 10.6 years, S.D.; PHQ-9 ≥ 10 or GAD-7 ≥ 10). All participants in the symptomatic group exhibited depression symptoms (PHQ-9 ≥ 10), with 64 of 85 showing comorbid anxiety symptoms (GAD-7 scores ≥ 10). Of all participants, 141 completed at least 8 stimulation sites (**Fig. S1A**). Detailed demographic statistics for each site are presented in **Table 1**, with statistics for two groups (healthy group and symptomatic group) separately provided in **Tables S1–S2**. All participants signed informed consent under ethical principles in the Declaration of Helsinki and all procedures were approved by the Stanford Institutional Review Board (31224).

### Data Collection Procedures

#### Visit 1: Magnetic resonance imaging

During the first scan day, structural MRI was collected from each participant using a 3T GE 750 scanner with an 8-channel head coil. The structural MRI was acquired using high resolution T1 weighted (T1w) (3D inversion SPGR) sequence: TR = 8.6 ms, TE = 3.4 ms, TI= 450 ms, flip angle = 15°, FOV = 22 cm, 184 slices, matrix 256 × 256, 1.0 × 0.9 × 0.9 mm acquisition resolution.

#### Stimulation sites

The TMS cortical sites included left/right frontal pole (L/R FP), left/right anterior middle frontal gyrus (L/R aMFG), left/right posterior middle frontal gyrus (L/R pMFG), right inferior frontal joint (R IFJ), right frontal eye field (R FEF), right pre-supplementary motor area (R preSMA), right primary motor cortex (R M1), and right inferior parietal lobule (R IPL) (**Fig. 1**). MNI coordinates for each site are listed in **Table 1**.

The coordinates of these stimulation sites were determined based on reports in previous studies. Specifically, the L/R aMFG (key nodes in ventral attention network), the L/R pMFG and R IPL (key nodes in frontoparietal control network), and the R FEF and R IFJ (key nodes in dorsal attention network) were defined according to coordinates identified in one of our previous TMS-fMRI studies (28), where an independent component analysis on a rs-fMRI dataset from 38 healthy individuals was performed to identify these cortical nodes in these networks. The L/R FP were localized based on the FP1 and FP2 EEG electrode locations in the international 10-20 system, which therefore varied slightly across participants. The averaged MNI coordinates were reported and presented in **Table 1** and **Fig. 1B**. The coordinates of R preSMA were from a prior study (29), and R M1 were positioned at the hand-knob region of the right (primary) motor cortex where a pulse of TMS induces a noticeable hand twitch. Greater coverage of the right hemisphere is due to overall scan time constraints. Note that for the R IPL site, the number of participants with valid data was fewer than the other sites (**Fig. 1C**). This was primarily because the R IPL target was introduced later in the study and was technically difficult to access when participants were lying down in the scanner.

#### Visit 2: Concurrent TMS-fMRI

Prior to the TMS-fMRI runs, neuronavigation was performed to localize each participant’s stimulation targets with the Visor2 software (ANT Neuro, Enschede, NL) outside the scanner room. Target coordinates were transformed from standard MNI space to the individual’s native space and stereotaxically identified on their T1w images loaded in the Visor 2. The corresponding scalp stimulation locations were then marked on a Lycra swim cap (Speedo USA) that participants wore in the scanner for TMS-fMRI runs.

Immediately before scanning, the individual’s resting motor threshold (MT) was defined in the MRI room as the minimal stimulation intensity that induced visually observable hand motor activity in at least 5 of 10 consecutive trials. The TMS intensity was set as 120% MT during the TMS-fMRI runs. If no obvious hand movement was observed within 10 minutes, stimulation was performed at 100% machine output. The distribution of stimulation intensity in all participants is presented in **Fig. S1B**.

During the TMS-fMRI runs, TMS pulses were delivered through an MRI-compatible air-cooled figure-eight TMS coil (MagPro MRI-B91), which was fixated by a custom-built holder and connected to a Magpro X100 stimulator (Magventure) in an adjoining room. Single pulses were presented with a jittered inter-stimulus interval ranging from 1–6 TRs to avoid the prediction of when TMS would be delivered. This rapid event-related design resulted in 68 stimulations per site over 167 volumes. For R M1 and R IPL, the TMS coil was perpendicular to the orientation of precentral gyrus (with the handle at angle of ca. 45° to the midline pointing in posterior direction) (64). For prefrontal sites, the TMS coil was oriented perpendicular to the orientation of middle frontal gyrus (with the handle at angle of ca. 90° to the midline) (65, 66). For all 11 TMS sites, the center of the TMS coil was positioned tangentially to the skull. The TMS coil was repositioned for each stimulation site by sliding the participant out of the scanner bore, adjusting the coil holder position, and returning the participant into the scanner bore. Before the first run, participants were informed that they could stop the current run or the entire experiment at any time if they could not tolerate the discomfort induced by TMS.

The TMS-fMRI images were acquired using the same scanner as used during Day 1, but with a TMS-compatible single-channel head coil that provides sufficient space for accommodating the TMS coil. The images were acquired using a T2w gradient-echo spiral in/out pulse sequence: TE = 30 ms, flip angle = 85°, FOV = 22 cm, pixel size = 3.4 mm, 31 axial slices (4 mm thick) covering the whole brain, slice spacing 1 mm, matrix 64 × 64, 2 excitations. A 300 ms break was set between volumes (TA = 2100 ms) to allow single-pulse TMS delivery without affecting functional volume acquisition as we have done in our previous work (28, 61), resulting in an effective TR of 2400 ms. An automated high-order shimming for spiral acquisitions was run prior to each TMS-fMRI run to reduce blurring and signal loss from field inhomogeneities. Heart rate (pulse oximeter clipped to the distal phalanx) and respiration (mid-thoracic strain gauge; both GE Healthcare) were acquired at 50 Hz during each fMRI run. They were processed offline and were regressed during data reconstruction, where raw P-files were converted to NIFTI files. During the scanning, participants wore earplugs to protect their hearing and minimize the impact of sounds from TMS clicks and scanner noise.

#### TMS-induced discomfort assessment

We recorded the Subjective Units of Distress Scale (SUDS) immediately after each TMS-fMRI run. SUDS was initially developed to measure anxiety during behavior therapy and has been widely used to index the current level of discomfort or distress experienced (27). Participants were asked to report the level of TMS-induced pain or discomfort from 0 to 100, with ‘0’ meaning ‘no pain’ and ‘100’ meaning ‘the worst pain you can imagine’.

### Data Analyses

#### Stimulation-site and group effects on TMS-induced discomfort

Following prior evidence that TMS-induced discomfort levels varied across stimulation locations (4) and mixed evidence regarding group differences in pain perception between healthy and clinical populations (54), we examined the SUDS ratings from different stimulation sites and two groups separately. To quantify the effects of stimulation site, group (healthy vs. symptomatic), and participant demographics (gender and age) on the level of TMS-induced discomfort feelings, we constructed a linear mixed-effects model:

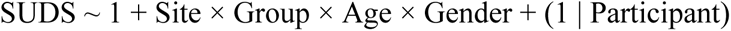

Participant identity was modeled as a random intercept to account for repeated measures and individual differences in the baseline of subjective reporting. Post-hoc comparisons were performed for any significant main effects or interactions.

#### MRI processing

Standard MRI preprocessing was performed using the fMRIPrep pipeline (67) as follows. The T1-weighted (T1w) image was processed with ANTs (68) for intensity nonuniformity correction, skull-stripping, and spatial normalization to the MNI template through nonlinear registration. Cerebrospinal fluid (CSF), white-matter (WM) and gray-matter (GM) were segmented on the brain-extracted T1w using FAST (FSL). For each of the TMS-fMRI run data, preprocessing steps included removal of the first 3 volumes, head-motion estimation using MCFLIRT (FSL), co-registration to the T1w reference using boundary-based registration with FLIRT (FSL), and resampling onto the standard MNI space.

Post-processing included 0.008 Hz high-pass filtering and 6-mm FWHM Gaussian kernel smoothing. Any fMRI runs with a maximum root mean square (RMS) absolute movement/displacement > 4 mm and/or mean RMS absolute movement/displacement > 2mm in any of the 6 motion parameters were excluded for quality control purposes. Next, a first-level general linear model (GLM) analysis was performed for each participant with SPM 12 (https://www.fil.ion.ucl.ac.uk/spm/). Individual TMS delivery timepoints were convolved with the canonical hemodynamic response function (HRF) with six head movement parameters as nuisance regressors. Following model estimation and contrast with implicit baseline, the parameter estimates (beta weights) were *z*-scored over whole-brain voxels and then averaged within 246 ROIs of the Brainnetome (BN) atlas (30) for subsequent analyses.

To assign each ROI to functional network(s), we first identified the Yeo 17-network (69) labels of all voxels within the ROI. The voxel-wise 17-network labels were then collapsed into 7 canonical networks, i.e., default mode, frontoparietal control, limbic, somatomotor, ventral attention, and visual networks. For each cortical ROI, we calculated the proportion of voxels assigned to each of these seven networks, and networks accounting for more than 20% of the ROI’s voxels were retained as the ROI’s functional network assignment. Subcortical ROIs were labeled separately as subcortical.

#### Modeling relationship between TMS-evoked brain responses and TMS-induced discomfort

We evaluated several multivariate computational frameworks that relate brain responses to SUDS ratings. The evaluated frameworks included the PCA-sCCA framework as established in our prior work (24, 25), as well as commonly used alternatives, including sCCA alone, sPLS alone, and the PCA-sPLS combination (26, 31). Model performance was compared to guide the selection of the primary computational framework for subsequent analyses.

#### PCA-sCCA framework

We first evaluated the PCA-sCCA computational framework. PCA is a commonly used dimensionality-reduction technique that transforms high-dimensional multivariate data into low-dimensional uncorrelated principal components (PCs). PCs explaining relatively low variance are assumed to primarily reflect noise and are therefore discarded (26). PCA was applied to the brain data, i.e., ROI-wise brain responses across all TMS-fMRI runs, to mitigate the potential multicollinearity problem within the brain data and improve the computational efficiency and robustness of the downstream computations. The transformation coefficients, i.e., the PCA loading matrix (246 × *k*), were obtained by matrix decomposition of the brain data. In our model, the number of retained PCs (*k*) was treated as a tunable hyperparameter and optimized in a data-driven manner. Following prior studies (25, 70) and considering the actual dimensionality of our dataset, the number of PCs was tested from 10 to 240 in increments of 10.

sCCA was then applied to the retained PCs to identify multivariate associations between TMS-evoked brain responses and TMS-induced discomfort. The sCCA is a multivariate latent variable model that identifies relationships between two data matrices (𝑿, 𝒀) by constructing canonical variates as linear combinations of variables within each matrix and maximizing the correlation between these variates (26). The coefficients of these linear combinations form the weight matrices (𝒘_𝒙_, 𝒘_𝒚_), which quantify the contribution of each variable within 𝑿 and 𝒀 to the association of the two matrices. In our model, the first data matrix 𝑿 comprises the *k* PCs derived from PCA; the second data matrix 𝒀 corresponds to the SUDS ratings. Because SUDS ratings are univariate, 𝒘_𝒙_ was a *k* × 1 vector and 𝒘_𝒚_was reduced to a scalar (set to 1). Sparsity was imposed on the 𝒘_𝒙_ via an L1-norm penalty to reduce model complexity, mitigate overfitting, and improve interpretability (70). The sparsity level was controlled by a hyperparameter *λ*, which was optimized in a data-driven manner. It was tested from 0.1 to 1 in increments of 0.1, with larger values yielding sparser solutions. To ensure comparable sparsity penalties across PCs, all PCs were variance-normalized (scaled to unit variance) (24). The SUDS ratings were standardized by *z*-scoring across samples. Here, the optimization problem to compute 𝒘_𝒙_was formulated as:

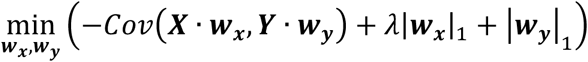

where 𝒘_𝒚_ = 1. The canonical variate for the brain responses was computed as 𝑿 · 𝒘_𝒙_, and its correlation *r* with 𝒀 quantified the strength of the brain-discomfort association. Given prior evidence that individuals with depressive or anxiety symptoms show marked alterations in brain systems involved in pain processing (20, 21), we separately modeled the healthy and symptomatic groups to identify population-specific brain-SUDS relationships.

#### Alternative multivariate frameworks

In addition to the PCA-sCCA framework, we evaluated several alternative frameworks, including sCCA alone, sPLS alone, and PCA-sPLS combination (26, 31). sPLS is a multivariate latent-variable model that constructs latent variates as linear combinations of variables (defined by sPLS weights 𝒘_𝒙_, *k* × 1) to maximize the covariance between the constructed variates (26). As in sCCA, sparsity was imposed on the sPLS weight vector for brain data using an L1-norm penalty. The sparsity hyperparameter was optimized in a data-driven manner and tested from 1 to sqrt(246) using logarithmically spaced sampling (26, 31).

For the sCCA-alone and sPLS-alone models, the ROI-wise brain responses (246 ROIs) served as the input without prior dimensionality reduction. For the PCA-sPLS model, PCA was first applied to the brain data, and the resulting PCs were subsequently entered into the sPLS model, with the number of retained PCs treated as a tunable hyperparameter using the same test range as in the PCA-sCCA framework.

#### Evaluation of model performance

Ten rounds of ten-fold cross-validation were performed to evaluate the generalizability and stability of the model and to guide hyperparameter optimization. During the cross-validation, in each fold, 90% of participants were assigned to the training set and the remaining 10% to the validation set (**Fig. S3**). All parameters estimated from the training set, e.g., the PCA loadings, sCCA weights, sPLS weights, etc., were applied to the validation set to derive the canonical or latent variates for the brain data. For each round, the derived variates from all ten folds were concatenated to form a full validation set. Pearson’s correlation coefficient (*r*_CV_) between the brain variates and SUDS ratings was computed to quantify model generalizability.

Model stability was assessed using the intraclass correlation coefficient (ICC) of the overall weight vector across cross-validation folds. For the PCA-sCCA model, the overall weight vector 𝑾 (246 × 1) was computed as the product of the PCA loading matrix (246 × *k*) and the sCCA weight vector (*k* × 1). Prior to this, the sCCA weight vector was first scaled by a variance-normalization vector (*k* × 1), defined as the standard deviations of the *k* retained PCs across samples. Each element of 𝑾 represents the contribution of a given ROI to the brain-discomfort association. To facilitate subsequent interpretation of the relative contribution, 𝑾 was further normalized by setting the largest absolute weight magnitude as 1.

For the PCA-sPLS model, the overall weight vector 𝑾 was computed analogously using the PCA loading matrix, the sPLS weight vector and the corresponding scaling vector. For the sCCA and sPLS alone models, 𝑾 was respectively computed as the element-wise product of the sCCA or sPLS weight vector (246 × 1) and a scaling vector (246 × 1), where the scaling vector consisted of the standard deviations of the TMS-evoked brain responses across the 246 ROIs.

Model performances across different frameworks were compared using *r*_CV_ and an additional combined metric that jointly incorporated *r*_CV_ and ICC (31). The combined metric was defined as one minus the Euclidean distance (ED) between the observed [*r*_CV_, ICC] and the ideal point [1, 1], which represents both perfect generalizability and stability. For both *r*_CV_ and the combined metric, higher values indicated better model performance. For each framework, hyperparameters were optimized to maximize the selected evaluation metric (24, 25).

The statistical significance of *r*_CV_ was further assessed using permutation testing (1,000 iterations). In each iteration, the correspondence between the TMS-evoked brain responses and SUDS ratings was randomly shuffled across samples, and the same cross-validation procedure was repeated to generate a null distribution of correlation coefficients. Statistical significance was determined by comparing the observed *r*_CV_ against this null distribution. All analyses were conducted separately for each group.

### Model selection and estimation of whole-brain weight patterns

For each group, we identified the optimal computational framework by comparing model performance across frameworks using both the *r*_CV_ and the combined metric. After selecting the optimal framework, *r*_CV_ was used to select the model hyperparameters (33). The final model was then re-applied to the full sample (i.e., without cross-validation) to estimate whole-brain weight patterns 𝑾 representing TMS-induced discomfort (**Fig. 3B**).

### Identification and statistical inference of key ROIs in the discomfort-related pattern

To facilitate interpretation, we focused on the top 15 ROIs with the largest absolute weight magnitudes in each group, following common criteria used in prior studies (25, 32, 33). Statistical significance of ROI-wise weights was assessed using permutation testing (10,000 iterations) by shuffling the correspondence between brain responses and SUDS ratings across samples. In each iteration, the optimal model was then applied to the permuted data to generate null distributions of ROI-wise weights. Resulting *p*-values were corrected for multiple comparisons across ROIs using the false discovery rate (FDR).

### Between-group comparison of discomfort-related ROI-wise weights

ROI-wise weights were compared between the groups for the selected top ROIs. The statistical significance of the observed between-group differences was then evaluated using the same permutation testing described above (10,000 iterations), in which a null distribution was obtained by contrasting the weights of the two groups estimated from the permuted data. Resulting *p*-values were FDR-corrected for multiple comparisons across ROIs.

### Characterizing the network distribution of key ROIs

The distribution of the seven functional networks among the top ROIs was quantified as the proportion of ROIs assigned to each network, calculated by dividing the ROI count for that network by the total number of key ROIs. If an ROI was assigned to n networks (n > 1), it contributed 1/n to the count of each of those networks.

### Quantifying discomfort-related proportion within TMS-evoked responses

To quantify the extent to which TMS-evoked brain responses reflected discomfort-related processes, we projected each sample’s TMS-evoked brain response vector onto the discomfort-related weight vector derived from the PCA-sCCA framework (**Fig. 5A**). For the *n*-th sample, the TMS-evoked brain response vector was denoted as 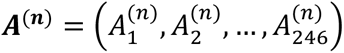 (corresponding to one row in the TMS-evoked response matrix 𝑨 shown in **Fig. 3**), where *a*_i_^(n)^ represents the response of the *i*-th ROI. The discomfort-related weight vector was denoted as 𝑾 = (𝑊_1_, 𝑊_2_, …, 𝑊_246._), where 𝑊𝑖 represents the weight for the 𝑖-th ROI. A least-squares regression model 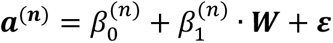 was used to estimate the discomfort-related projection ***d****^(n)^* = 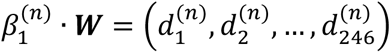, where *d*_i_^(n)^ represents the discomfort-related projection of the *i*-th ROI. The regression coefficient *β*_i_^(n)^ was estimated as:

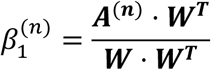

For the *i*-th ROI in the *n*-th sample, the proportion of the discomfort-related projection within the TMS-evoked response was computed as 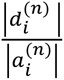. At the group level, we quantified the average of the discomfort-related proportion across samples as:

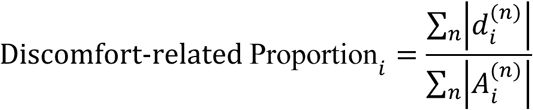

### Evaluating the group specificity of the discomfort-related neural signature

For each group, we first evaluated the brain-discomfort association in the full sample using the optimal model and hyperparameters. We then removed the discomfort-related projection 𝒅^(𝒏)^from the TMS-evoked brain responses for each sample and re-evaluated the correlation between the SUDS ratings and the brain canonical variate derived from the residual brain responses. Two complementary analyses were conducted. In the within-group analysis, 𝒅^(𝒏)^ was computed using the discomfort-related weight patterns derived from the same group; in the cross-group analysis, 𝒅^(𝒏)^was computed using weight patterns derived from the other group. Statistical significance of the resulting correlations was assessed using permutation testing (10,000 iterations). In each iteration, the correspondence between the residual brain responses and SUDS ratings was randomly shuffled across samples, and the same PCA-sCCA model with optimized hyperparameters was reapplied to the permuted data to generate a null distribution for statistical testing. Correlations after projection removal were further compared with those obtained from the analysis prior to projection removal, with the reduction in correlation strength quantifying the contribution of the removed weight pattern to the brain-discomfort association.

## Acknowledgement

This work was supported by National Institutes of Health R01MH136197 (J.J.), R01MH129694 (Y.Z.), R21AG080425 (Y.Z.), R01MH103324 (A.E.), R01MH132074 (A.D.B.), R01NS114405 (A.D.B.), R01MH139650 (A.D.B.), R61MH135428 (D.J.O.), R01MH132740 (D.J.O.), R01MH129694 (D.J.O.), Brain and Behavior Research Foundation Young Investigator grant 29441 (J.J.), Roy J. Carver Trust (A.D.B.), and the Hart Fund in Cognitive Neuroscience (D.J.O.).

## Author contributions

Conceptualization: Z.L., J.J.; Writing – original draft: Z.L., J.J.; Writing – review and editing: Z.L., Y.J., Y.Z., N.Z., A.E., A.D.B., D.J.O., J.J.; Investigation: Z.L., J.J.; Formal analysis: Z.L., Y.J., J.J.; Methodology: Z.L., Y.J., N.Z., J.J.; Funding acquisition, resources, and supervision: Y.Z., A.E., A.D.B., D.J.O., J.J.

