## Supplementary information for "Whole-brain mapping of TMS-induced discomfort with large-scale concurrent TMS-fMRI"

**Figure S1. Distribution of participants by number of TMS sites completed and stimulation intensity.**
(A) TMS sites. Most participants completed ≥8 stimulation sites (healthy: 85.0%, left; symptomatic: 84.7%, right).
(B) TMS intensity. Most participants received stimulation at ≥90% of maximum stimulator output (healthy: 88.8%, left; symptomatic: 95.3%, right).

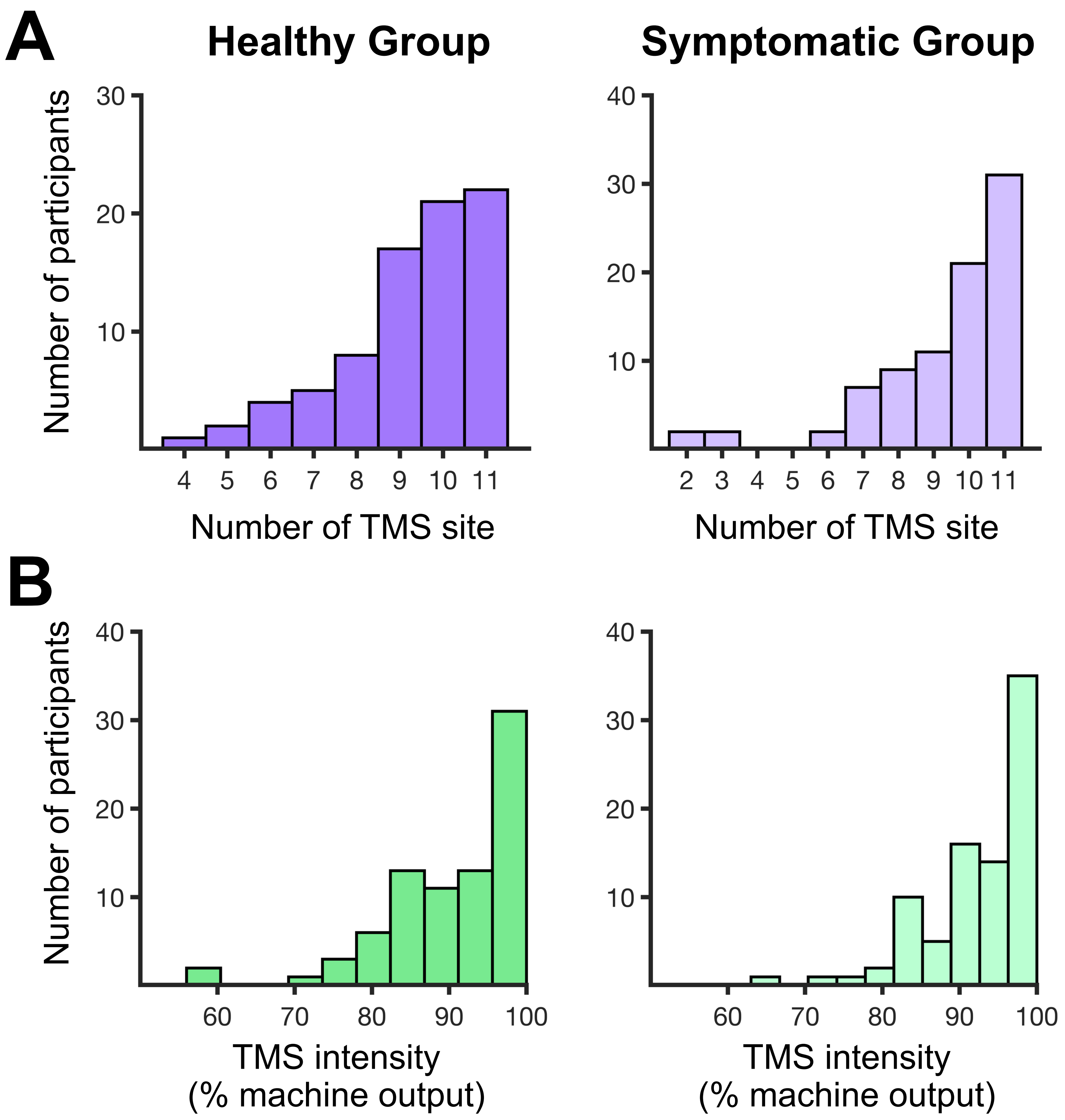

**Figure S2. TMS-induced discomfort across stimulation sites for the two groups.**
Mean SUDS ratings at each stimulation site and pairwise cross-site comparisons in the healthy group (A) and in the symptomatic group (B). Statistically significant differences are marked by asterisks ($p_{FDR} < 0.05$), and marginally significant differences by crosses ($p_{FDR} < 0.10$). Overall, higher discomfort ratings were observed at more frontal stimulation sites.

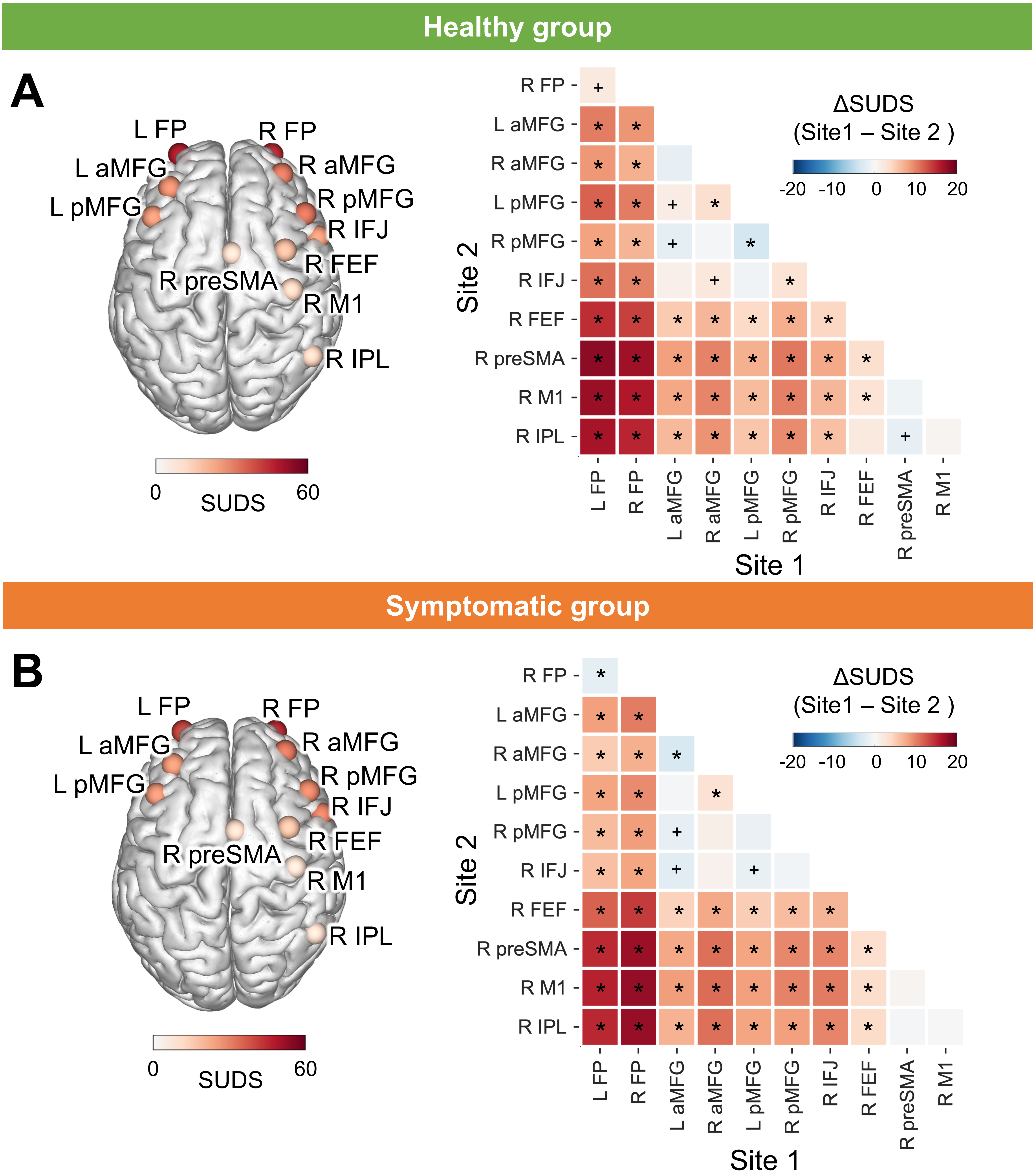

**Figure S3. Cross-validation strategy and optimal hyperparameters of the PCA-sCCA framework.**
(A) Cross-validation strategy for the PCA-sCCA computational framework. Ten rounds of ten-fold cross-validation was performed to evaluate model performance. In each fold, 90% of participants were assigned to the training set and 10% to the validation set. All parameters estimated from the training set, including PCA loadings and sCCA weights, were applied to the validation set to generate the cross-validated canonical variate for the brain responses. The correlation $r_{CV}$ between the brain canonical variate and the SUDS ratings of the validation set was computed for quantifying the cross-validated generalizability of model (see Methods for details).
(B-C) Optimal hyperparameters by maximizing $r_{CV}$ for the healthy (B) and symptomatic groups (C). The optimal model retained 160 PCs with the sparsity parameter of $\lambda$ = 0.25 for the healthy group and 50 PCs with $\lambda$ = 0.20 for the symptomatic group.

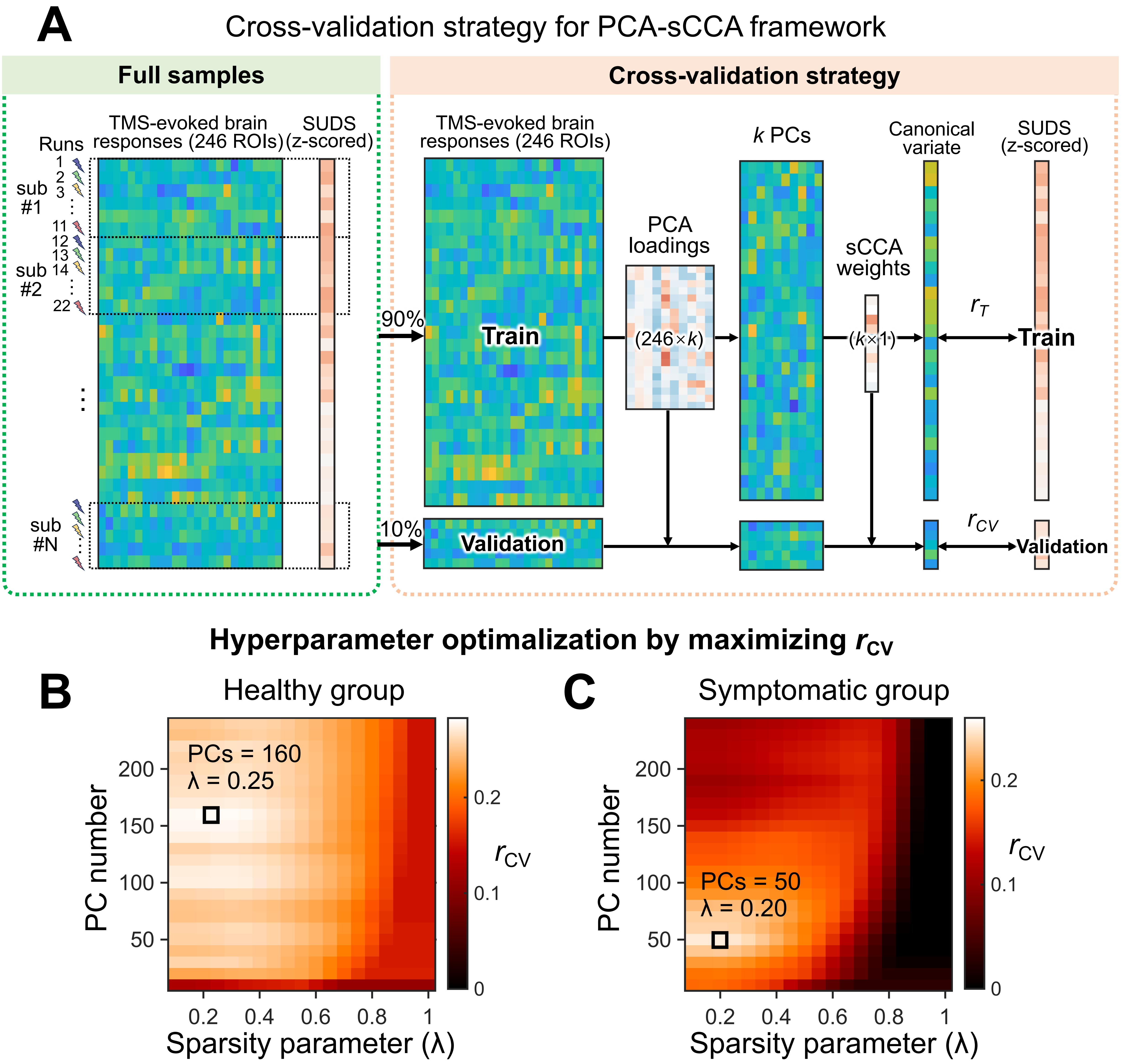

**Figure S4. Cross-validated model generalizability with the optimal hyperparameters.**
For both the healthy group (A-B) and symptomatic group (C-D), all ten rounds of cross-validation yielded statistically significant brain-discomfort associations ($p_{\text{permutation}} < .001$).

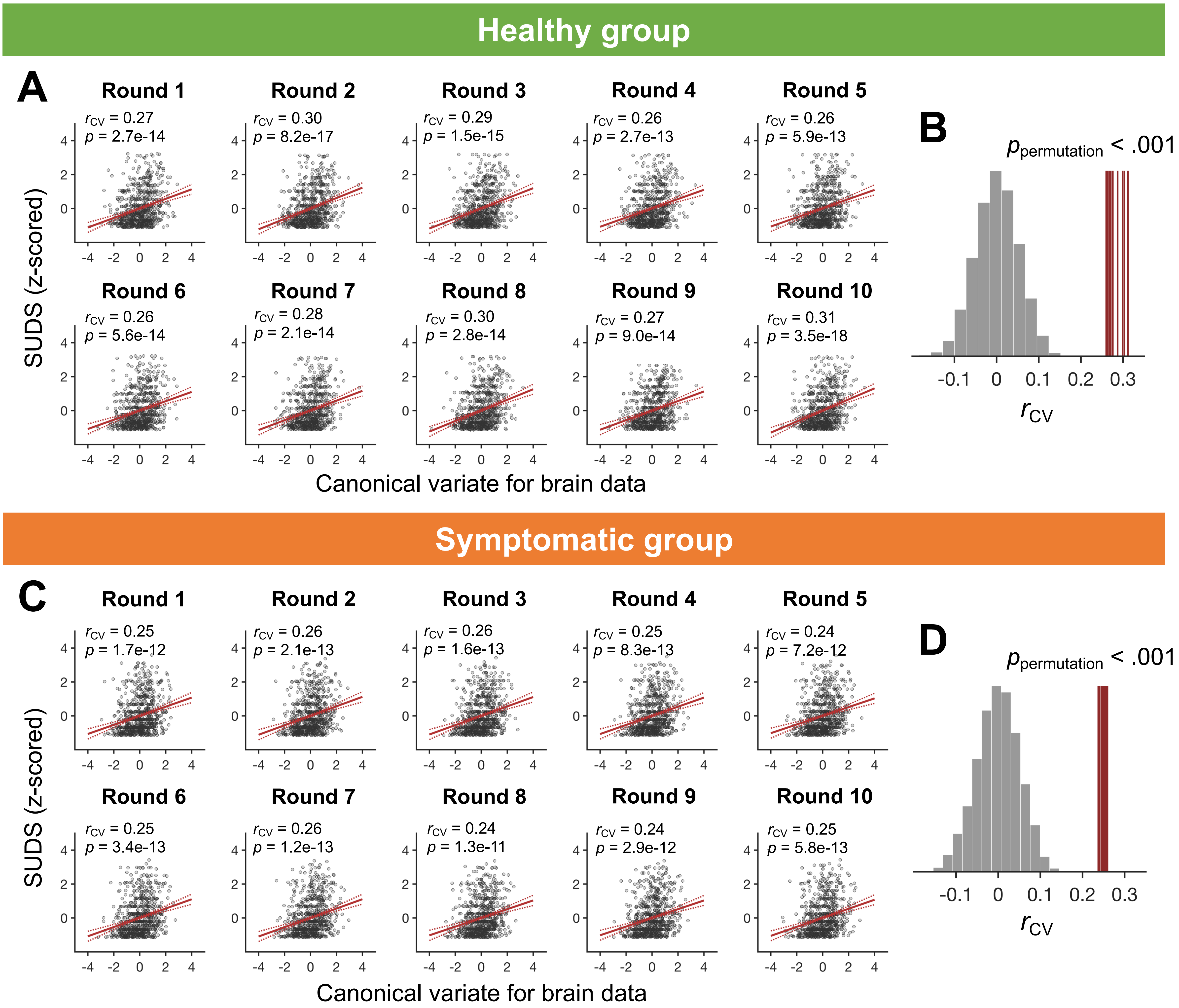

**Figure S5. PCA-sCCA framework outperformed alternative multivariate frameworks.**
We evaluated and compared the performance of several multivariate frameworks to our primary framework (PCA-sCCA), including sCCA alone, sPLS alone, and PCA-sPLS (see Methods). The same cross-validation strategy as in the PCA-sCCA framework was used to evaluate the model performance. The cross-validated correlation $r_{CV}$ and a combined metric that jointly incorporated $r_{CV}$ and model stability, quantified by the intraclass correlation coefficient (ICC), were used to compare the model performance. For both metrics, higher values indicate better performance.
(A–C) Results in the healthy group. (A) Performance of different multivariate frameworks across hyperparameter configurations in the healthy group. Each marker represents one hyperparameter configuration, and markers with different colors and shapes indicate different frameworks. (B–C) Maximum $r_{CV}$ (B) and combined metric (C) for each frameworks, with the optimal hyperparameters identified for maximizing the corresponding metrics and for each framework. The PCA-sCCA framework shows the highest performance across both metrics.
(D–F) Results in the symptomatic group. The PCA-sCCA framework shows the best performance in both $r_{CV}$ and the combined metric.

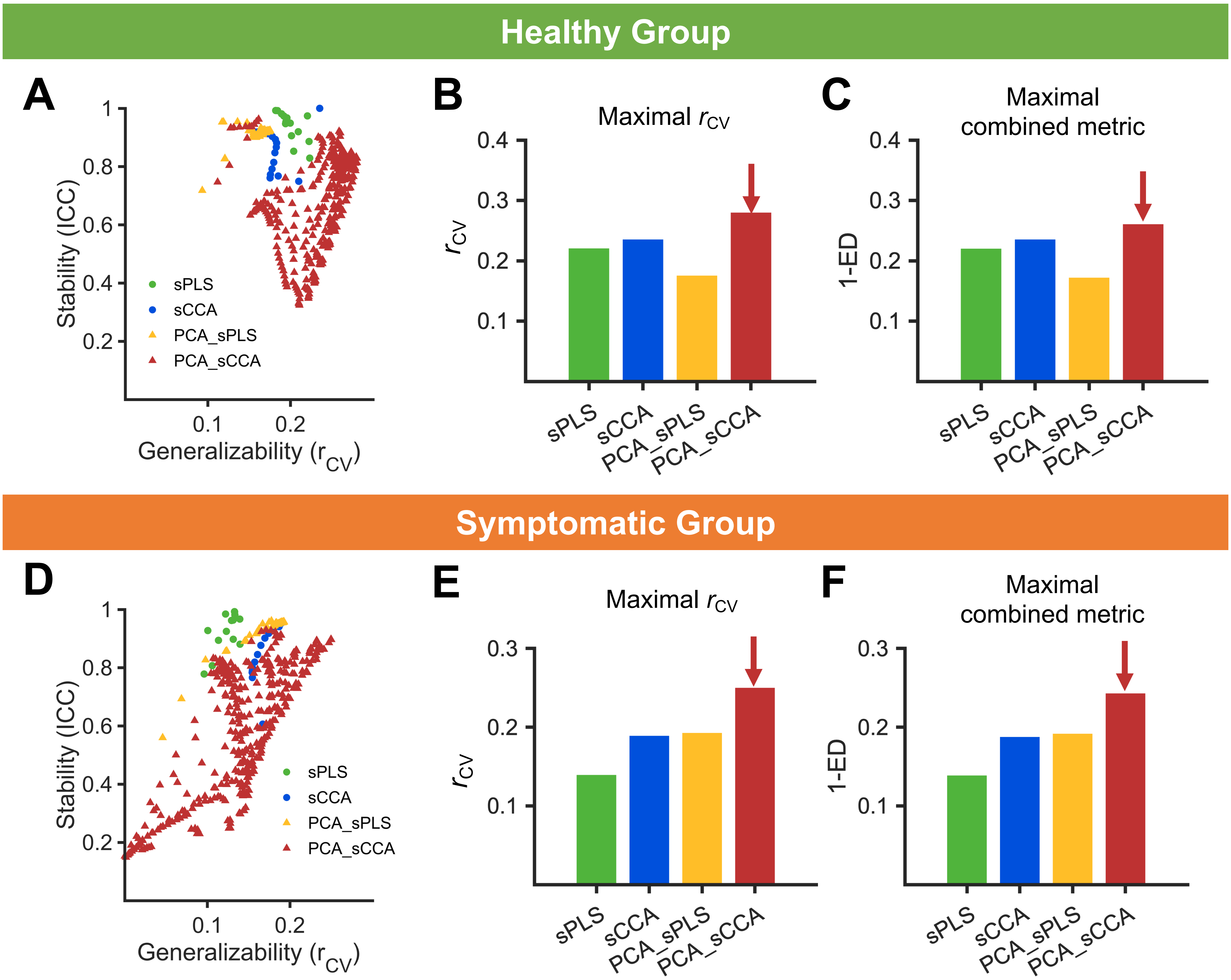

**Figure S6. Significance of weights in the top 15 ROIs.**
For both the healthy group (A) and symptomatic group (B), the weights of top 15 ROIs exhibited statistically significant ($p_{FDR} < 0.05$) or marginally significant ($p_{FDR} < 0.07$) by permutation testing (N = 10,000; see Methods).

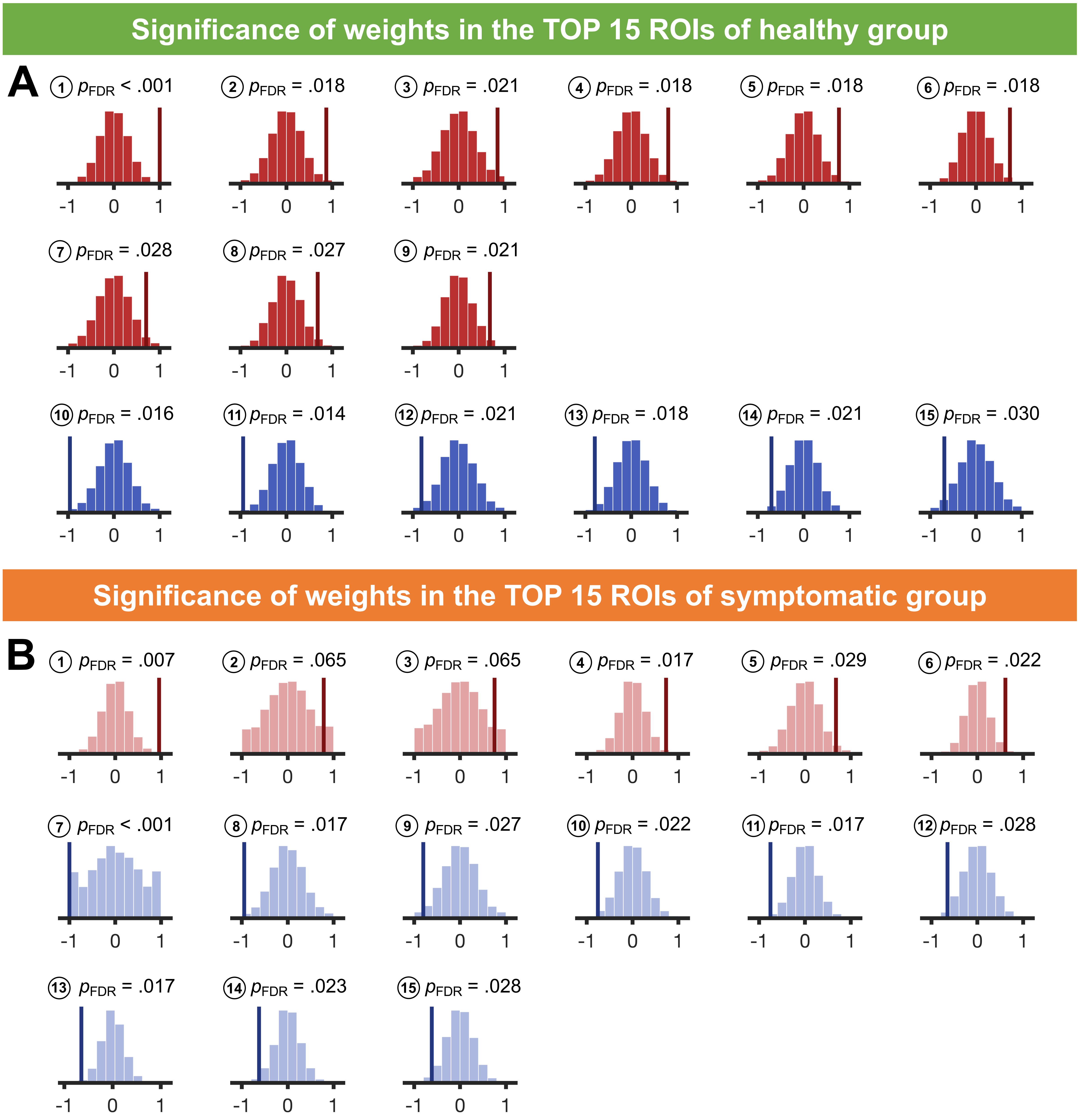

**Figure S7. Association between SUDS ratings and TMS-evoked brain responses before and after regressing out the whole-brain discomfort-related pattern.**

(A) Healthy group. After regressing out the whole-brain weight pattern identified from the healthy group (middle), the correlation was significantly reduced relative to that from the raw data (left) ($z$ = 6.06, $p$ = 6.68e–10) and was no longer statistically significant ($p_{permutation}$ = .90). In contrast, removing the weight pattern identified from the symptomatic group minimally affected the correlation, which remained significant ($p_{permutation}$ < .001) (right).

(B) Symptomatic group. After regressing out the symptomatic-specific weight pattern (middle), the correlation was significantly reduced ($z$ = 3.85, $p$ = 5.87e–5) and no longer significant ($p_{permutation}$ = .36). Regressing out the healthy-group pattern had little impact, and the correlation remained significant (right).

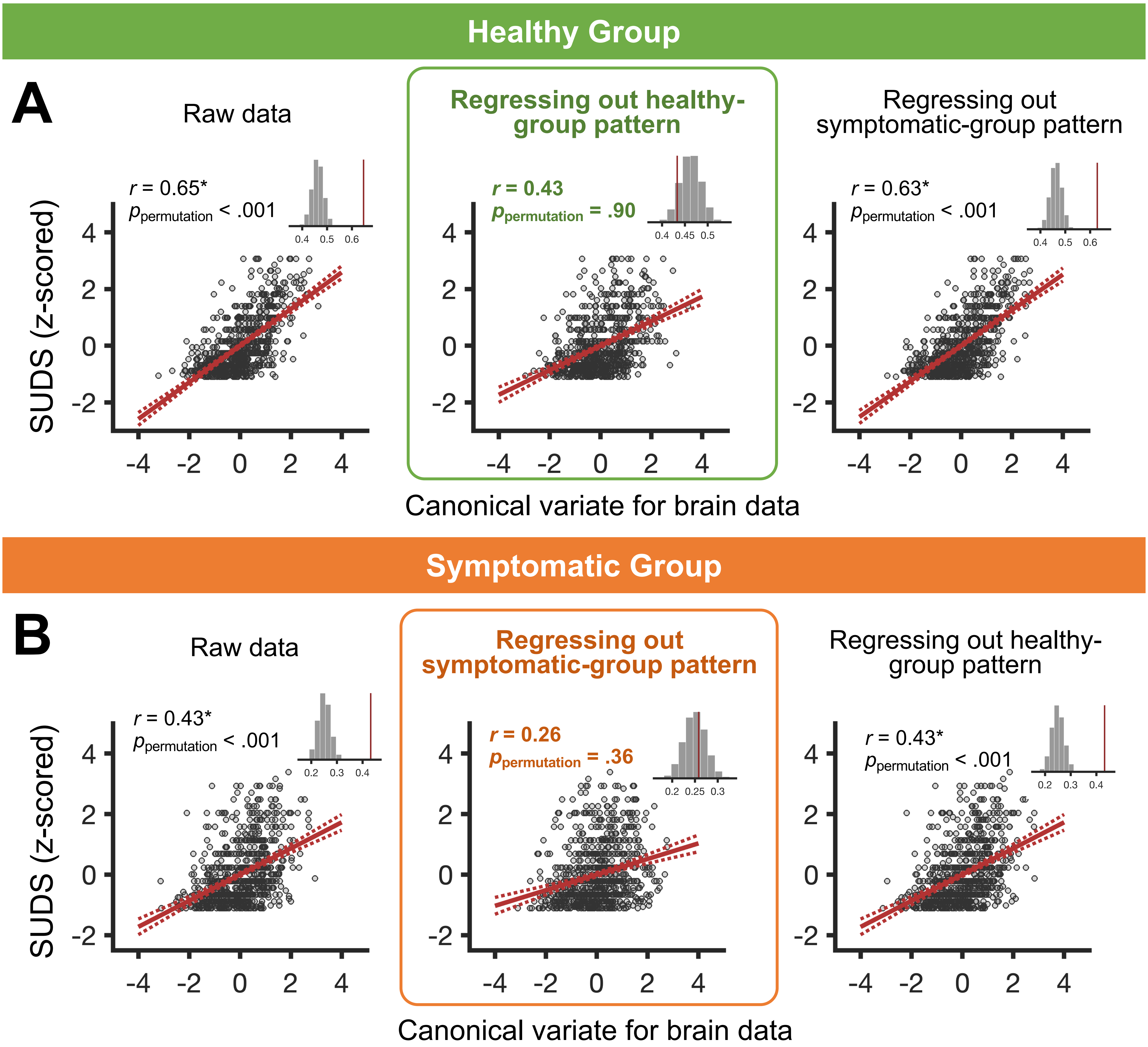

**Table S1. Number of participants, demographic information, and MNI coordinates for each TMS site in the healthy group.**

| TMS sites | Number of participants | Female | Male | Age(S.D.) | Years of Education (S.D.) | MNI coordinates | | |
|---|---|---|---|---|---|---|---|---|
| | | | | | | x | y | z |
| L Fp | 63 | 36 | 27 | 32.3(11.3) | 16.2(2.2) | -24 | 60 | -2 |
| R Fp | 62 | 34 | 28 | 33.2(11.2) | 16.4(2.2) | 24 | 60 | -2 |
| L aMFG | 75 | 44 | 31 | 30.8(10.2) | 16.0(2.1) | -32 | 42 | 34 |
| R aMFG | 74 | 43 | 31 | 31.8(10.6) | 16.2(2.1) | 30 | 50 | 26 |
| L pMFG | 77 | 45 | 32 | 31.7(10.6) | 16.1(2.2) | -38 | 22 | 38 |
| R pMFG | 78 | 46 | 32 | 31.5(10.6) | 16.1(2.1) | 46 | 26 | 38 |
| R IFJ | 71 | 43 | 28 | 31.3(10.9) | 16.1(2.2) | 46 | 8 | 28 |
| R FEF | 73 | 46 | 27 | 31.7(10.9) | 16.0(2.1) | 34 | 6 | 62 |
| R preSMA | 58 | 34 | 24 | 30.8(10.6) | 16.1(2.1) | 6 | 2 | 68 |
| R M1 | 73 | 44 | 29 | 32.0(10.6) | 16.2(2.0) | 40 | -18 | 64 |
| R IPL | 38 | 23 | 15 | 32.0(11.5) | 16.4(2.2) | 48 | -54 | 46 |
| **All** | 80 | 47 | 33 | 31.7(10.6) | 16.1(2.1) | - | - | - |

**Table S2. Number of participants, demographic information, and MNI coordinates for each TMS site in the symptomatic group.**

| TMS sites | Number of participants | Female | Male | Age(S.D.) | Years of Education (S.D.) | MNI coordinates | | |
|---|---|---|---|---|---|---|---|---|
| | | | | | | x | y | z |
| L Fp | 66 | 42 | 24 | 31.2(10.9) | 15.5(1.8) | -24 | 60 | -2 |
| R Fp | 66 | 42 | 24 | 31.1(10.9) | 15.6(1.9) | 24 | 60 | -2 |
| L aMFG | 75 | 49 | 26 | 31.1(10.4) | 15.6(1.9) | -32 | 42 | 34 |
| R aMFG | 77 | 53 | 24 | 31.0(10.6) | 15.7(1.8) | 30 | 50 | 26 |
| L pMFG | 78 | 52 | 26 | 31.4(10.8) | 15.6(1.9) | -38 | 22 | 38 |
| R pMFG | 80 | 54 | 26 | 31.3(10.7) | 15.7(1.9) | 46 | 26 | 38 |
| R IFJ | 74 | 49 | 25 | 32.0(10.8) | 15.5(1.9) | 46 | 8 | 28 |
| R FEF | 80 | 54 | 26 | 30.9(10.4) | 15.7(1.9) | 34 | 6 | 62 |
| R preSMA | 73 | 48 | 25 | 30.1(9.8) | 15.6(2.0) | 6 | 2 | 68 |
| R M1 | 76 | 51 | 25 | 31.0(10.6) | 15.6(1.9) | 40 | -18 | 64 |
| R IPL | 48 | 31 | 17 | 27.6(7.7) | 15.3(1.8) | 48 | -54 | 46 |
| **All** | 85 | 57 | 28 | 31.0(10.5) | 15.6(1.9) | - | - | - |

**Table S3. Substantial interindividual variability in TMS-induced discomfort within each stimulation site.** IQR = interquartile range; Q1 = first quartile; Q3 = third quartile; S.D. = standard deviation.

| TMS site | Both group | | | Healthy group | | | Symptomatic group | | |
|---|---|---|---|---|---|---|---|---|---|
| | **Mean (S.D.)** | **Range [min, max]** | **IQR [Q1,Q3]** | **Mean (S.D.)** | **Range [min, max]** | **IQR [Q1,Q3]** | **Mean (S.D.)** | **Range [min, max]** | **IQR [Q1,Q3]** |
| L FP | 43.2(27.0) | [5, 100] | [20, 66] | 46.6(30.1) | [5, 100] | [20, 73] | 39.9(23.4) | [6, 90] | [20, 55] |
| R FP | 44.3(25.6) | [1, 100] | [20, 65] | 44.2(27.9) | [1, 100] | [20, 68] | 44.4(23.5) | [1, 100] | [25, 60] |
| L aMFG | 26.4(20.1) | [0, 90] | [10, 40] | 27.5(19.6) | [1, 85] | [10, 42] | 25.3(20.7) | [0, 90] | [10, 35] |
| R aMFG | 31.4(23.5) | [0, 100] | [11, 45] | 31.3(24.3) | [0, 100] | [15, 40] | 31.5(22.9) | [0, 90] | [10, 50] |
| L pMFG | 25.6(20.3) | [0, 90] | [10, 37] | 25.3(20.6) | [0, 80] | [10, 38] | 25.9(20.0) | [0, 90] | [10, 35] |
| R pMFG | 30.4(23.1) | [0, 95] | [10, 45] | 32.1(24.2) | [0, 90] | [10, 50] | 28.7(22.0) | [0, 95] | [10, 40] |
| R IFJ | 27.4(20.1) | [0, 80] | [10, 40] | 25.2(19.3) | [0, 70] | [10, 40] | 29.6(20.6) | [0, 80] | [13, 40] |
| R FEF | 17.4(18.1) | [0, 80] | [5, 25] | 18.1(19.4) | [0, 75] | [5, 21] | 16.7(17.0) | [0, 80] | [5, 25] |
| R preSMA | 10.4(13.3) | [0, 70] | [2, 10] | 10.0(13.0) | [0, 60] | [2, 10] | 10.7(13.6) | [0, 70] | [2, 11] |
| R M1 | 10.7(13.1) | [0, 70] | [2, 14] | 12.3(14.3) | [0, 50] | [2, 16] | 9.1(11.7) | [0, 70] | [2, 10] |
| R IPL | 10.6(11.0) | [0, 50] | [3, 15] | 12.4(12.0) | [0, 50] | [5, 16] | 9.2(10.1) | [0, 40] | [2, 12] |

**Table S4. Optimal model performance of various multivariate frameworks.**
Cross-validated generalizability $r_{CV}$ and a combined metric jointly incorporating $r_{CV}$ and model stability, quantified by the intraclass correlation coefficient (ICC), were computed to evaluate and compare frameworks. The composite metric was defined as one minus the Euclidean distance (ED) between the [$r_{CV}$, ICC] and the ideal point [1, 1] (see Methods). For both $r_{CV}$ and the composite metric, higher values mean better model performances. For each framework, optimal hyperparameters were selected to maximize the corresponding evaluation metric. Across both groups and for both metrics, PCA-sCCA exhibited the best model performance among the four frameworks..

| Group | | Maximize $r_{CV}$ | | | Maximize 1-ED | | |
|---|---|---|---|---|---|---|---|
| | | $r_{CV}$ | ICC | 1-ED | $r_{CV}$ | ICC | 1-ED |
| Healthy | sPLS | 0.22 | 0.83 | 0.20 | 0.22 | 0.97 | 0.22 |
| | sCCA | 0.24 | 1.00 | 0.24 | 0.24 | 1.00 | 0.24 |
| | PCA-sPLS | 0.18 | 0.92 | 0.17 | 0.18 | 0.92 | 0.17 |
| | **PCA-sCCA** | **0.28** | 0.83 | **0.26** | **0.28** | 0.83 | **0.26** |
| Symptomatic | sPLS | 0.14 | 0.88 | 0.13 | 0.14 | 0.97 | 0.14 |
| | sCCA | 0.19 | 0.95 | 0.19 | 0.19 | 0.95 | 0.19 |
| | PCA-sPLS | 0.19 | 0.96 | 0.19 | 0.19 | 0.96 | 0.19 |
| | **PCA-sCCA** | **0.25** | 0.90 | **0.24** | **0.25** | 0.90 | **0.24** |

**Table S5. Rank and statistical significance of weights for top ROIs in the healthy group**

For each ROI, the Brainnetome (BN) atlas index, architectonic parcellation, network assignment, weight, weight rank among all 246 ROIs, and statistical significance assessed using permutation testing (10,000 iterations) are reported.

| | BN index | ROI | Network | Weights | Rank | $p_{perm}$ | $p_{FDR}$ |
|---|---|---|---|---|---|---|---|
| 1 | 58 | R sPreCG | SMN | 1.00 | 1 | <0.0001 | <0.001 |
| 2 | 62 | R iPreCG | SMN, VAN | 0.87 | 4 | 0.006 | 0.018 |
| 3 | 157 | L iPoCG(PO, S2) | SMN | 0.85 | 5 | 0.017 | 0.021 |
| 4 | 41 | L mPFC | DMN | 0.81 | 7 | 0.010 | 0.018 |
| 5 | 189 | L cLG | VN | 0.77 | 9 | 0.010 | 0.018 |
| 6 | 191 | L rCun | VN | 0.74 | 10 | 0.008 | 0.018 |
| 7 | 227 | L dCN | SC | 0.70 | 11 | 0.026 | 0.028 |
| 8 | 180 | R dACC | VAN | 0.68 | 14 | 0.024 | 0.027 |
| 9 | 96 | R mITG | LN, DMN | 0.68 | 15 | 0.016 | 0.021 |
| | **BN index** | **ROI** | **Network** | **Weights** | **Rank** | $p_{perm}$ | $p_{FDR}$ |
| 10 | 188 | R pgACC | DMN | -0.96 | 2 | 0.003 | 0.016 |
| 11 | 67 | L aPCL | SMN | -0.94 | 3 | 0.002 | 0.014 |
| 12 | 219 | L vCN | SC | -0.81 | 6 | 0.014 | 0.021 |
| 13 | 25 | L PMv | FPN, DAN | -0.79 | 8 | 0.009 | 0.018 |
| 14 | 172 | R dGI | SMN, VAN | -0.70 | 12 | 0.014 | 0.021 |
| 15 | 48 | R mOFC | LN | -0.69 | 13 | 0.030 | 0.030 |

**Table S6. Rank and statistical significance of weights for top ROIs in the symptomatic group**

| | BN index | ROI | Network | Weights | Rank | $p_{perm}$ | $p_{FDR}$ |
|---|---|---|---|---|---|---|---|
| 1 | 145 | L SMG | SMN, VAN | 0.96 | 2 | 0.001 | 0.007 |
| 2 | 166 | R vAI | VAN | 0.79 | 5 | 0.064 | 0.065 |
| 3 | 115 | L mPHG | LN, DMN | 0.75 | 8 | 0.065 | 0.065 |
| 4 | 157 | L iPoCG(PO, S2) | SMN | 0.73 | 9 | 0.007 | 0.017 |
| 5 | 170 | R vDI/GI | VAN | 0.68 | 10 | 0.025 | 0.029 |
| 6 | 158 | R iPoCG(PO, S2) | SMN | 0.61 | 15 | 0.012 | 0.022 |
| | **BN index** | **ROI** | **Network** | **Weights** | **Rank** | $p_{perm}$ | $p_{FDR}$ |
| 7 | 117 | L aPHG | LN | -1.00 | 1 | <0.0001 | <0.001 |
| 8 | 114 | R pPHG | DMN | -0.95 | 3 | 0.005 | 0.017 |
| 9 | 77 | L aSTG | LN, DMN | -0.80 | 4 | 0.018 | 0.027 |
| 10 | 219 | L vCN | SC | -0.76 | 6 | 0.011 | 0.022 |
| 11 | 155 | L mPoCG (S1) | SMN | -0.75 | 7 | 0.006 | 0.017 |
| 12 | 113 | L pPHG | DMN | -0.65 | 11 | 0.020 | 0.028 |
| 13 | 156 | R mPoCG (S1) | SMN | -0.65 | 12 | 0.004 | 0.017 |
| 14 | 20 | R aMFG | FPN, VAN | -0.62 | 13 | 0.014 | 0.023 |
| 15 | 228 | R dCN | SC | -0.62 | 14 | 0.023 | 0.028 |

Abbreviations: aMFG = anterior middle frontal gyrus; aPCL = anterior paracentral lobule; aPHG = anterior parahippocampal gyrus; aSTG = anterior superior temporal gyrus; cLG = caudal lingual gyrus; dACC = dorsal anterior cingulate cortex; dCN = dorsal caudate nucleus; dGI = dorsal granular insula; iPoCG = inferior postcentral gyrus (parietal operculum, PO; secondary somatosensory cortex, S2); iPreCG = inferior precentral gyrus; mITG = middle inferior temporal gyrus; mOFC = medial orbitofrontal cortex; mPFC = medial prefrontal cortex; mPHG = middle parahippocampal gyrus; mPoCG = middle postcentral gyrus (primary somatosensory cortex, S1); pPHG = posterior parahippocampal gyrus; pgACC = pregenual anterior cingulate cortex; PMv = ventral premotor cortex; SMG = supramarginal gyrus; rCun = rostral cuneus; sPreCG = superior precentral gyrus; vAI = ventral anterior insula; vCN = ventral caudate nucleus; vDI/GI = ventral dysgranular/granular insula.

**Table S7. Between-group comparison of the weights for top ROIs in the healthy group.**
For each ROI, the weight in the healthy group ($w_H$), the weight in the symptomatic group ($w_S$), the between-group weight difference ($w_H$ - $w_S$), and the statistical significance of the group difference assessed using permutation testing (10,000 iterations) are reported. ROIs showing statistically significant group differences ($p_{FDR} < 0.05$) are highlighted in red.

| | ROI | Network | $w_H$ | $w_S$ | $w_H - w_S$ | $p_{perm}$ | $p_{FDR}$ |
|---|---|---|---|---|---|---|---|
| 1 | **R sPreCG** | SMN | **1.00** | -0.20 | **1.20*** | 0.001 | 0.015 |
| 2 | R iPreCG | SMN, VAN | **0.87** | 0.33 | **0.54** | 0.099 | 0.185 |
| 3 | L iPoCG(PO, S2) | SMN | **0.85** | 0.73 | **0.11** | 0.407 | 0.422 |
| 4 | L mPFC | DMN | **0.81** | 0.37 | **0.44** | 0.165 | 0.243 |
| 5 | L cLG | VN | **0.77** | 0.32 | **0.45** | 0.154 | 0.240 |
| 6 | L rCun | VN | **0.74** | 0.33 | **0.41** | 0.133 | 0.218 |
| 7 | L dCN | SC | **0.70** | -0.45 | **1.16*** | 0.007 | 0.031 |
| 8 | **R dACC** | VAN | **0.68** | 0.13 | **0.55** | 0.110 | 0.192 |
| 9 | R mITG | LN, DMN | **0.68** | 0.35 | **0.33** | 0.186 | 0.261 |
| | **ROI** | **Network** | $w_H$ | $w_S$ | $w_H - w_S$ | $p_{perm}$ | $p_{FDR}$ |
| 10 | **R pgACC** | DMN | **-0.96** | 0.46 | **-1.42*** | 0.001 | 0.015 |
| 11 | L aPCL | SMN | **-0.94** | -0.25 | -0.69 | 0.044 | 0.103 |
| 12 | L vCN | SC | **-0.81** | -0.76 | -0.06 | 0.456 | 0.456 |
| 13 | **L PMv** | FPN, DAN | **-0.79** | 0.24 | **-1.03*** | 0.005 | 0.031 |
| 14 | **R dGI** | SMN, VAN | **-0.70** | 0.38 | **-1.08*** | 0.006 | 0.031 |
| 15 | **R mOFC** | LN | **-0.69** | 0.36 | **-1.05*** | 0.009 | 0.031 |

**Table S8. Between-group comparison of the weights for top ROIs in the symptomatic group.**
ROIs showing statistically significant ($p_{FDR} < 0.05$) or marginally significant ($p_{FDR} < 0.07$) group differences are highlighted in red.

| | ROI | Network | $w_S$ | $w_H$ | $w_S - w_H$ | $p_{perm}$ | $p_{FDR}$ |
|---|---|---|---|---|---|---|---|
| 1 | **L SMG** | SMN, VAN | **0.96** | -0.26 | **1.22*** | 0.004 | 0.031 |
| 2 | R vAI | VAN | **0.79** | 0.40 | **0.39** | 0.264 | 0.284 |
| 3 | L mPHG | LN, DMN | **0.75** | -0.06 | **0.81** | 0.082 | 0.165 |
| 4 | L iPoCG(PO, S2) | SMN | **0.73** | 0.85 | **-0.11** | 0.407 | 0.422 |
| 5 | **R vDI/GI** | VAN | **0.68** | -0.26 | **$0.94^{+}$** | 0.025 | 0.063 |
| 6 | R iPoCG(PO, S2) | SMN | **0.61** | 0.30 | **0.32** | 0.233 | 0.283 |
| | **ROI** | **Network** | $w_S$ | $w_H$ | $w_S - w_H$ | $p_{perm}$ | $p_{FDR}$ |
| 7 | **L aPHG** | LN | **-1.00** | 0.25 | **$-1.25^{+}$** | 0.018 | 0.056 |
| 8 | R pPHG | DMN | **-0.95** | -0.59 | **-0.36** | 0.244 | 0.284 |
| 9 | **L aSTG** | LN, DMN | **-0.80** | 0.41 | **-1.21*** | 0.009 | 0.031 |
| 10 | L vCN | SC | **-0.76** | -0.81 | **0.06** | 0.456 | 0.456 |
| 11 | L mPoCG (S1) | SMN | **-0.75** | -0.53 | **-0.23** | 0.263 | 0.284 |
| 12 | L pPHG | DMN | **-0.65** | -0.29 | **-0.36** | 0.224 | 0.283 |
| 13 | **R mPoCG (S1)** | SMN | **-0.65** | -0.01 | **$-0.63^{+}$** | 0.021 | 0.059 |
| 14 | R aMFG | FPN, VAN | **-0.62** | 0.02 | **-0.64** | 0.073 | 0.157 |
| 15 | R dCN | SC | **-0.62** | -0.27 | **-0.35** | 0.227 | 0.283 |

**Table S9. Proportion of discomfort-related responses in the TMS-evoked responses across stimulation sites and groups.**

For each TMS site, we quantified the proportion of discomfort-related responses within the TMS-evoked brain responses. Mean (S.D.) values were computed across ROIs, separately for the top 15 ROIs with the largest weight magnitude, for the remaining ROIs (excluding top 15 ROIs) and for all 246 ROIs.

| | Healthy group | | | Symptomatic group | | |
|---|---|---|---|---|---|---|
| **TMS sites** | **Top 15 ROIs** | **Remaining ROIs** | **All ROIs** | **Top 15 ROIs** | **Remaining ROIs** | **All ROIs** |
| L FP | 17.2(3.9)% | 5.5(4.3)% | 6.2(5.1)% | 30.6(6.9)% | 10.1(7.2)% | 11.4(8.7)% |
| R FP | 17.8(3.8)% | 5.5(4.2)% | 6.2(5.1)% | 32.9(7.0)% | 11.1(7.8)% | 12.4(9.3)% |
| L aMFG | 11.1(2.5)% | 3.3(2.6)% | 3.8(3.2)% | 23.4(5.7)% | 7.7(5.4)% | 8.7(6.6)% |
| R aMFG | 14.3(2.8)% | 4.3(3.3)% | 4.9(4.1)% | 26.7(5.6)% | 8.8(6.2)% | 9.9(7.5)% |
| L pMFG | 12.1(3.0)% | 3.5(2.8)% | 4.0(3.5)% | 24.8(5.6)% | 8.3(5.8)% | 9.3(7.0)% |
| R pMFG | 13.0(2.5)% | 4.0(3.0)% | 4.5(3.7)% | 26.8(6.3)% | 8.9(6.3)% | 10.0(7.6)% |
| R IFJ | 11.6(2.4)% | 3.5(2.7)% | 4.0(3.3)% | 27.6(6.3)% | 8.6(6.1)% | 9.7(7.6)% |
| R FEF | 11.4(2.7)% | 3.3(2.5)% | 3.8(3.2)% | 23.8(5.3)% | 7.7(5.4)% | 8.7(6.6)% |
| R preSMA | 9.4(1.9)% | 2.7(2.1)% | 3.1(2.6)% | 21.8(5.4)% | 6.9(4.9)% | 7.8(6.1)% |
| R M1 | 7.6(1.5)% | 2.3(1.8)% | 2.6(2.2)% | 19.8(3.3)% | 6.2(4.3)% | 7.0(5.4)% |
| R IPL | 9.9(2.2)% | 3.0(2.3)% | 3.4(2.8)% | 19.9(3.7)% | 6.1(4.4)% | 6.9(5.5)% |
| **All** | **12.3(2.4)%** | **3.7(2.8)%** | **4.2(3.5)%** | **25.2(4.9)%** | **8.2(5.7)%** | **9.2(7.0)%** |
